# Saturation mutagenesis of a predicted ancestral Syk-family kinase

**DOI:** 10.1101/2022.04.24.489292

**Authors:** Helen T. Hobbs, Neel H. Shah, Sophie R. Shoemaker, Jeanine F. Amacher, Susan Marqusee, John Kuriyan

## Abstract

Many tyrosine kinases cannot be expressed readily in *E. coli*, limiting facile production of these proteins for biochemical experiments. We used ancestral sequence reconstruction to generate a spleen tyrosine kinase (Syk) variant that can be expressed in bacteria and purified in soluble form, unlike the human members of this family (Syk and ZAP-70). The catalytic activity, substrate specificity, and regulation by phosphorylation of this Syk variant are similar to the corresponding properties of human Syk and ZAP-70. Taking advantage of the ability to express this novel Syk-family kinase in bacteria, we developed a two-hybrid assay that couples the growth of *E*.*coli* in the presence of an antibiotic to successful phosphorylation of a bait peptide by the kinase. Using this assay, we screened a site-saturation mutagenesis library of the kinase domain of this reconstructed Syk-family kinase. Sites of loss-of-function mutations identified in the screen correlate well with residues established previously as critical to function and/or structure in protein kinases. We also identified activating mutations in the regulatory hydrophobic spine and activation loop, which are within key motifs involved in kinase regulation. Strikingly, one mutation in an ancestral Syk-family variant increases the soluble expression of the protein by 75-fold. Thus, through ancestral sequence reconstruction followed by deep mutational scanning, we have generated Syk-family kinase variants that can be expressed in bacteria with very high yield.

## 1. INTRODUCTION

Eukaryotic protein kinases catalyze the transfer of a phosphoryl group from ATP to serine, threonine, or tyrosine residues in their substrate proteins.^1,2^ The timing and location of such phosphorylation is essential for the regulation of many cellular processes, such as growth, development, and responses to extracellular stimuli. Thus, the dysregulation of protein kinases frequently results in disease, including cancer and auto-immune disorders.^3^ Decades of work have revealed the mechanisms of regulation, the molecular structures, and the substrate specificities of most kinases.^4,5^ However, many kinases do not express well in bacteria, which limits the throughput of structural and functional studies.^6^ For some kinases, the co-expression of a phosphatase enables kinase expression, suggesting that kinase activity is toxic to bacteria.^6^ This strategy is not universally successful,^7^ most likely because these eukaryotic proteins do not fold efficiently in bacteria. Strategies to overcome this problem include the systematic mutagenesis of solvent exposed hydrophobic residues and the co-expression of chaperones.^8–11^

Here, we show that ancestral sequence reconstruction can be used to identify tyrosine kinase variants that can be expressed and purified in soluble form from bacteria. In this method, ancestral protein sequences are inferred from phylogenetic trees and multiple sequence alignments encompassing a large set of evolutionarily related proteins.^12^ We applied ancestral sequence reconstruction to the spleen tyrosine kinase (Syk) family, which is comprised of two members, Syk and zeta-chain-associated protein kinase of 70 kDa (ZAP-70).^13^ Syk and ZAP-70 have the same domain architecture, in which a tandem SH2 module is followed by a tyrosine kinase domain (Figure 1A and B), and they play corresponding roles in B cells and T cells, respectively.^14^ Previous work characterizing their structure and function has been carried out using protein expressed in, and purified from, SF21 insect cells, as neither human Syk nor human ZAP-70 can be expressed in bacteria, even with co-expression of a tyrosine phosphatase (Figure 1C).^15,16^ We show that the predicted common ancestor of Syk and ZAP-70 expresses robustly as soluble protein in *E. coli*. Several previous studies have demonstrated that the proteins predicted by ancestral sequence reconstruction are more thermostable than their currently-existing counterparts,^17,18^ which may allow such proteins to fold more readily in bacteria.

**Figure 1.**
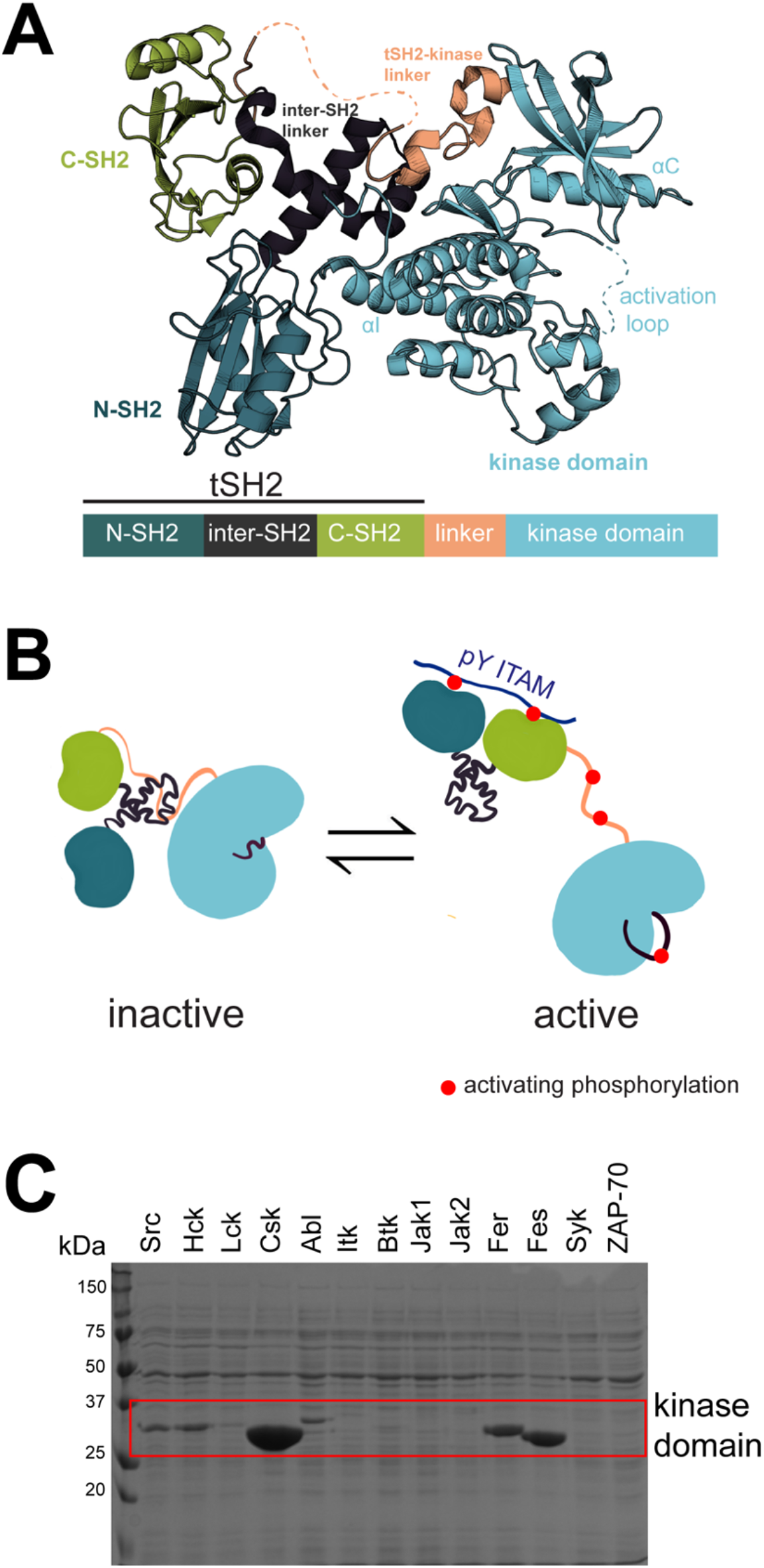
The Syk-family kinases. **A**. The structure of full-length auto-inhibited human ZAP-70 kinase (PDB: 4K2R) **B**. Activation of the Syk-family kinases by a conformational change in the tSH2 module and phosphorylation of key tyrosine residues (indicated with red circles). The tSH2 module binds to a doubly-phosphorylated immunoreceptor tyrosine-based activation motif (ITAM) and phosphorylation of tSH2-kinase linker stabilizes the open, active conformation. The activation loop is also phosphorylated for maximal activation. **C**. His-tagged human kinase domains expressed in *E. coli* co-expressing the YopH phosphatase (MW=45kDa) and enriched over nickel columns. The expected molecular weight of each tyrosine kinase domain falls within the red box (between 25 and 37 kDa). A strong band in this box indicates soluble expression. Human ZAP-70 and Syk, the last two lanes, show no soluble expression in *E. coli* with YopH.

Using the bacterially-expressed ancestral Syk kinase, we developed a bacterial two-hybrid assay for tyrosine kinase activity based on the interaction between a phosphotyrosine-containing peptide and a Src-homology (SH2) domain. This assay enabled a saturation mutational scan, in which every residue in the kinase domain of the ancestral Syk-family kinase was replaced by all 19 other amino acid residues, one at a time. Deep-mutagenesis experiments rely on high-throughput functional assays, in which many genetically-encoded protein variants are sequenced before and after functional screening or selection. Previous high-throughput selection assays for kinase activity have relied on yeast^19,20^ or mammalian cells.^19,21–24^ While powerful, these approaches can be challenging, due to the slow growth rate of these cells, native kinase signaling, and technical difficulties in screening large libraries. Bacteria, on the other hand, grow rapidly, can be transformed easily with large plasmid libraries, and do not have any canonical protein tyrosine kinases. Additionally, previous work has demonstrated that bacterial screens can be used to reveal new insights into the allosteric regulation and stability of eukaryotic signaling proteins.^25,26^

The saturation-mutagenesis scan of the bacterially-expressed Syk family kinase revealed that the two-hybrid assay can identify residues that result in loss-of-function when mutated, and that these residues have been established previously as critical for function and/or structure in protein tyrosine kinases. Purification of variants with activating mutations identified a variant of the predicted ancestral Syk kinase that is expressed at even higher levels (∼75-fold greater) than the original ancestral Syk kinase. Thus, through two steps, ancestral sequence reconstruction followed by saturation mutagenesis, we have identified a Syk-family kinase with extremely high expression in bacteria (∼150 mgs of purified protein from 1 L of bacterial culture). Bacterially-expressed variants of difficult to express kinases, such as Syk and ZAP-70, could serve as useful models for studying the activity, regulation, specificity, and inhibition of a diverse set of biologically important kinases.

## 2. RESULTS AND DISCUSSION

### 2.1 Ancestral sequence reconstruction yields a Syk-family kinase that can be expressed in *E. coli*

To carry out the ancestral sequence reconstruction, we first made a multiple sequence alignment of 183 Syk-family kinases found in present-day metazoan species, ranging from sponges to primates (Supplementary File 1).^27^ This alignment was used to generate a phylogenetic tree (Figure 2A and Supplementary File 2), the topology of which reflected taxonomic relationships between the relevant species^28^ and was consistent with a known gene duplication event in this lineage that occurred with the emergence of jawed vertebrates.^29^ The multiple sequence alignment and phylogenetic tree were used as the input for ancestral sequence reconstruction using the program Lazurus.^30^ This process resulted in the reconstruction of ancestral Syk-family kinases corresponding to internal nodes in the phylogenetic tree. We chose three of these nodes for further study, corresponding to a common ancestor of Syk and ZAP-70 (AncSZ), a Syk ancestor (AncS), and a ZAP-70 ancestor (AncZ). The pairwise sequence identities of the reconstructed and human Syk-family kinases are shown in Figure 2B, for both the full-length proteins and the kinase domains. The kinase domain of the common ancestor, AncSZ, is 71% identical to the kinase domain of human ZAP-70 and 74% identical to the kinase domain of human Syk. A sequence alignment of the kinase domains of human ZAP-70, AncZ, AncSZ, AncS and human Syk can be found in Figure 2C.

**Figure 2.**
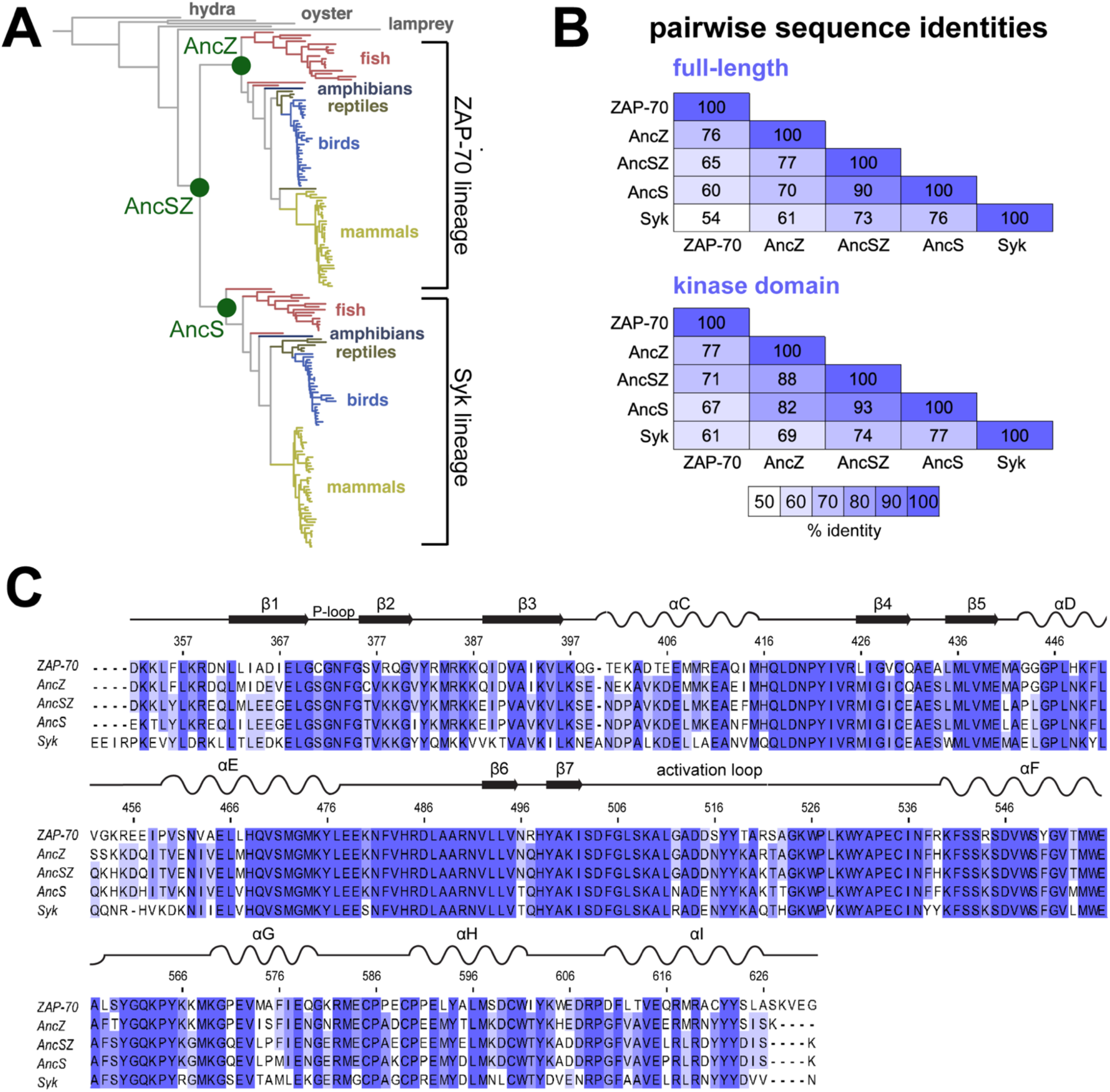
Reconstructed Syk-family kinases. **A**. The phylogenetric tree used in the ancestral sequence reconstruction. The nodes corrresponding to AncS, AncZ, and AncSZ are marked with green circles. **B**. The pairwise sequence identities of the full-length and kinase domains of human ZAP-70, AncZ, AncSZ, AncS, and human Syk. **C**. A sequence alignment of the kinase domains of the human and reconstructed Syk-family kinases. Residues are shaded according to percent identity. Numbered according to AncSZ.

We attempted to express AncSZ, AncZ, and AncS in Sf21 insect cells as well as in *E. coli* Bl21(DE3) cells co-expressing the tyrosine phosphatase YopH.^6^ As for human Syk and ZAP-70, AncS and AncZ could only be expressed in insect cells. The common ancestor, AncSZ, could, however, be expressed in soluble form as both the full-length construct (∼0.1 mg of purified protein obtained from 1 liter of *E. coli* culture) and the isolated kinase domain (∼2 mg/L yield) (Figure 3A). Expression of these proteins required co-expression of a tyrosine phosphatase, YopH.^6^ Strikingly, saturation mutagenesis of AncSZ, discussed in more detail below, identified a mutation, L616R (corresponding to Gln 591 in ZAP-70 and Leu 624 in Syk), that further increased the expression of soluble protein, by approximately 75-fold (with a yield of ∼150 mg of soluble purified protein per liter of *E. coli* culture; Figure 3A). The identification of this super-expressing Syk kinase variant, dubbed AncSZ*, demonstrates that a combination of ancestral sequence reconstruction and saturation mutagenesis can be used to generate a protein tyrosine kinase variant with extremely high expression of soluble protein in bacteria.

**Figure 3.**
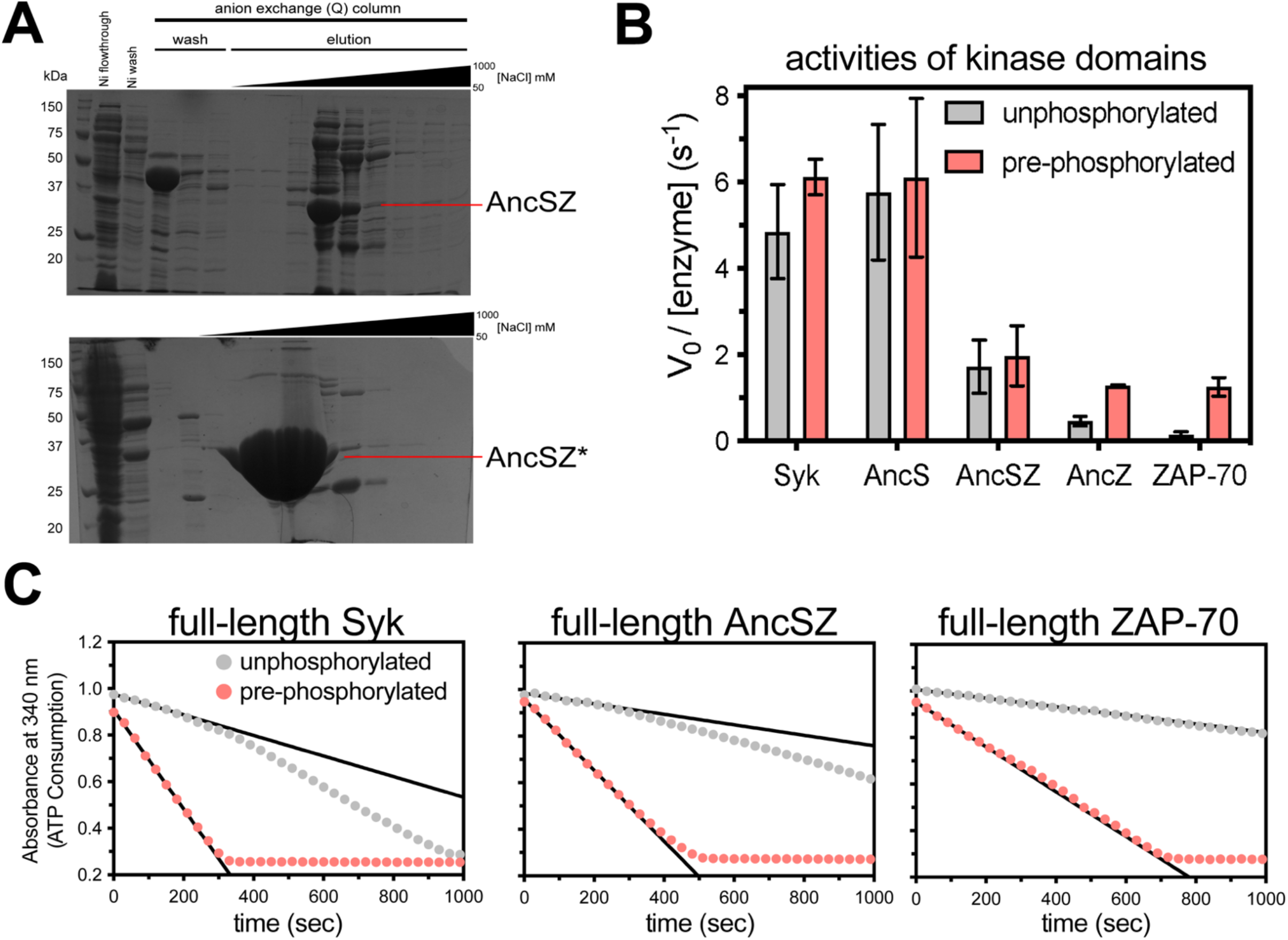
Characterization of the bacterially-expressed AncSZ and other Syk-family kinases. **A**. Top, the protein purification gel of AncSZ when expressed in *E. coli* co-expresssing the protein tyrosine phosphatase YopH. The band corresponding to YopH (45kDa) is observed in the lanes corresponding to the anion exchange (Q) column washes. A band at the expected molecular weight of AncSZ (∼32kDa) is observed in fractions taken during the elution of AncSZ from the Q column with increasing concentration of NaCl. The yield from this purification was approximately 2 mg of protein from one liter of *E. coli* culture. Bottom, a similar gel from the purification of AncSZ*. The yield for this protein was approximately 150 mg of protein from one liter of *E. coli* culture, a substantial increase over what was observed for AncSZ. **B**. The catalytic activities (the initial velocities) of ancestral and extant Syk-family kinases against LAT_214-233_ (n=3). For the pre-phosphorylated samples, kinases were incubated with purified Lck kinase and ATP for one hour prior to measurement. **C**. Reaction progress curves for full-length Syk, ZAP-70, and AncSZ phosphorylation of LAT_214-233_, measured in an enzymatic assay where ADP production is coupled to NADH oxidation and a loss of absorbance at 340 nm. The black lines in each graph track the initial velocities. Both Syk and AncSZ can auto-activate, as indicated by the increasing reaction rate for the unphosphorylated samples as a function of time. ZAP-70, however, cannot.

### 2.2 Reconstructed ancestral Syk-family kinases have biochemical properties characteristic of human Syk and ZAP-70

We compared the catalytic activities of the bacterially-expressed and purified AncSZ protein with those of the kinases expressed in SF21 cells, including human ZAP-70, human Syk, AncS, and AncZ. The catalytic activities of the purified kinase domains were measured with and without prior phosphorylation of the activation loop (Figure 3B and Figure SI 1A). Phosphorylation of this loop stabilizes the active conformations of most kinases.^31^ Activation-loop phosphorylation was achieved via pre-incubation with a Src-family kinase, Lck, which is an endogenous regulator of ZAP-70 in T cells.^32^ The addition of Lck is necessary because activation-loop auto-phosphorylation is slow for both the human and ancestral kinases and is especially so for ZAP-70 and AncZ, compared to Syk and AncS, consistent with previous reports (Figure SI 2).^33^ As expected, the Lck-phosphorylated forms of the ZAP-70 and AncZ kinase domains were significantly more active than unphosphorylated samples (Figure 3B). For Syk and AncS, activation loop phosphorylation by Lck had a negligible effect on activity, as previously observed for Syk.^15^ The common ancestor, AncSZ, had an intermediate catalytic activity, higher than that of ZAP-70 and AncZ, but lower than the activities of Syk and AncS. As for Syk and AncS, AncSZ activity in the isolated kinase domain was not significantly impacted by Lck-mediated activation loop phosphorylation (Figure 3B).

We were also able to express and purify full-length AncSZ from *E. coli*. Full-length Syk-family kinases adopt an auto-inhibited conformation when unphosphorylated, and are activated by phosphorylation of the linker connecting the tandem SH2 domains to the kinase domain as well as by phosphorylation of the activation loop (Figure 1B).^34,35^ We measured the catalytic activities toward a peptide substrate with and without pre-phosphorylation by Lck of full-length AncSZ, purified from *E. coli*, and full-length ZAP-70 and Syk, purified from insect cells (Figure SI 1B). As for full-length human Syk and ZAP-70, the activity of the bacterially-expressed full-length AncSZ was substantially higher when pre-phosphorylated by Lck (Figure 3C). As seen for the isolated kinase domains, we found that Syk was the most active kinase and ZAP-70 was least active, with AncSZ falling in between (Figure 3C). In the samples without Lck, we observed a slow increase in catalytic efficiency for Syk, but not for ZAP-70, consistent with the ability of Syk to auto-phosphorylate its SH2-kinase linker (Figure 3C).^15,35^ AncSZ was also able to auto-activate, albeit more slowly than Syk.

The restricted substrate specificity of the human Syk-family kinases is an important feature of their biological function, and substrates that are phosphorylated efficiently by Syk-family kinases are usually poor substrates for Src-family kinases.^33,36^ To assess the substrate specificities of the reconstructed Syk-family kinases, we used the kinase domains to phosphorylate a library of ∼3000 diverse peptides that are tyrosine kinase substrates, using a previously described bacterial surface-display assay (Figure 4A).^33,36,37^ Briefly, the peptide library is expressed in *E. coli*, and phosphorylation of the surface-displayed peptides is detected by the binding of fluorescently-labeled antibody recognizing phosphorylated tyrosine residues. Cells bearing highly phosphorylated peptides are sorted using flow cytometry, and the peptide-encoding DNA is sequenced. Comparison with the input population of peptide-encoding DNA variants provides a measure of the efficiency with which each peptide is phosphorylated.^33,36^

**Figure 4.**
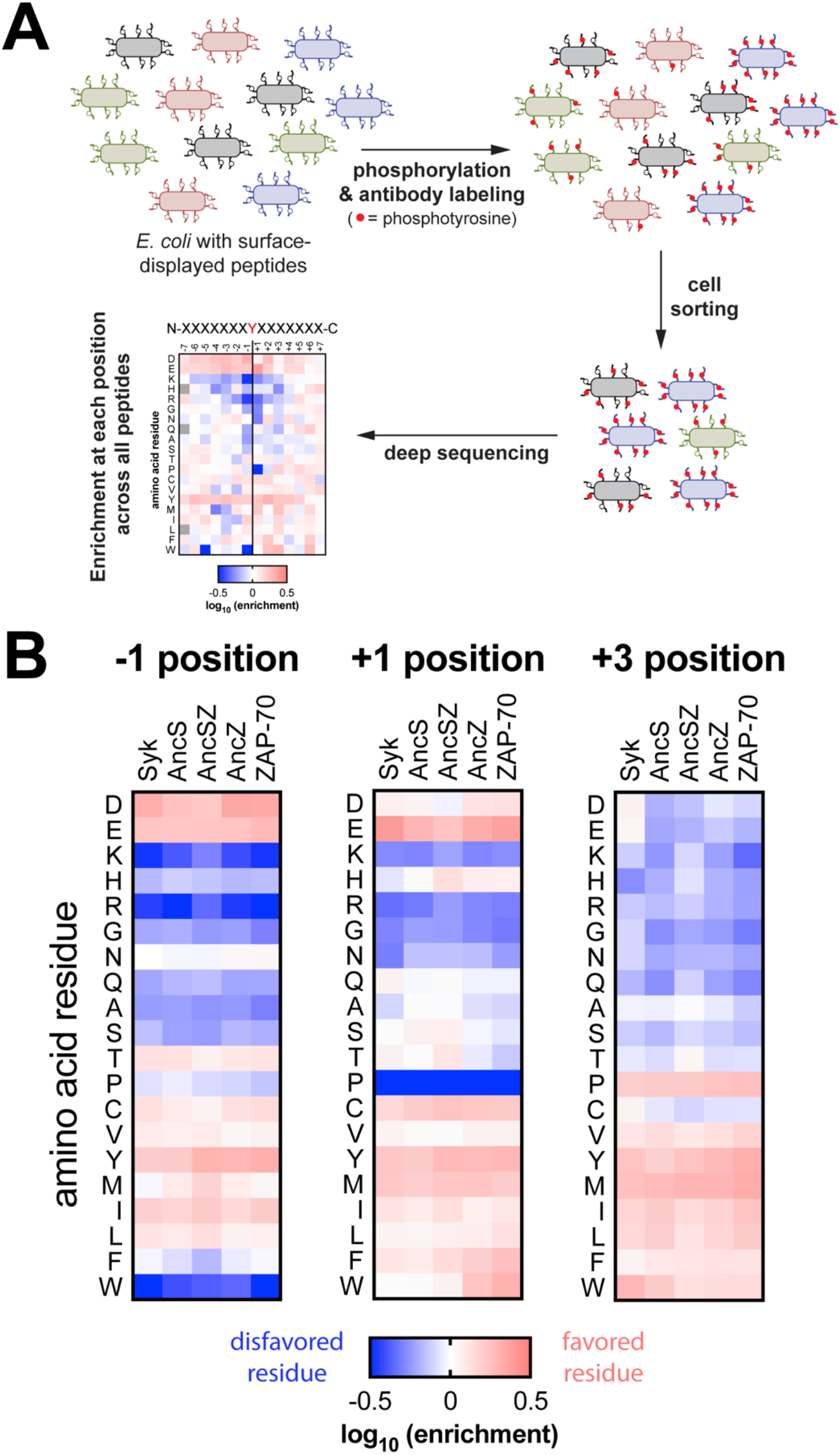
The predicted ancestral Syk-family kinases retain the substrate specificity profile of the human Syk-family kinases. **A**. *E. coli* are transformed with a plasmid library of ∼3000 diverse peptides fused to a bacterial surface-display scaffold. Displayed peptides are then phosphorylated by the addition of the kinase variant of interest and labeled with a fluorescent pan-phosphotyrosine antibody. Labeled cells are sorted according to fluorescence and peptide-encoding plasmids from cells in the selected and unselected populations are isolated for deep sequencing, allowing for the calculation of enrichment at each position in the peptide. **B**. The enrichment of amino acids at positions (−1, +1, +3) in the substrate that are known determinants of substrate specificity in Syk-family kinases.

The reconstructed ancestral Syk kinases retain the substrate-specificity patterns that are characteristic of Syk-family kinases (Figure 4B and Figure SI 3), of which the most important is a bias against positively-charged residues across the substrate peptide. Like the human Syk-family kinases, the ancestral kinases exhibit a strong preference for an acidic residue at the position before the phosphotyrosine (denoted -1) but will also tolerate a bulky aliphatic amino acid (L/I) (Figure 4B). The ancestral kinases also demonstrate a slight preference for glutamate at the +1 position and a proline at the +3 position, as has been reported for the human proteins.^33,38,39^, again recapitulating the known substrate specificity for present-day Syk-family kinases.

### 2.3 Construction of a high-throughput assay for the kinase activity of AncSZ

We designed and implemented a bacterial two-hybrid assay for tyrosine kinase activity in *E. coli* (Figure 5A). In a typical two-hybrid assay, expression of a reporter gene (e.g. an antibiotic resistance marker) is coupled to the successful interaction between a “bait” and “prey” protein. Here, we add a third component, the kinase, that must first phosphorylate the bait, a tyrosine-containing peptide, in order for it to interact with the prey, a Src-homology 2 (SH2) domain. The components of the assay are expressed on three plasmids and are adapted from a previously reported bacterial two-hybrid assay for protein-protein interactions.^25,40^ The bait, a 20-residue peptide spanning Tyr 226 in the ZAP-70 substrate LAT, is fused to the λ-cI protein by a (Gly-Ser)_3_ linker and is expressed from a pZS22 plasmid (Figure SI 4A). The prey protein corresponds to the Src-homology 2 (SH2) domain of the adapter protein Grb2 (residues 60-152), which binds to phosphorylated LAT Tyr 226.^41^ The Grb2 SH2 domain is fused to the N-terminal domain of the α-subunit of *E. coli* RNA polymerase via a flexible (Gly-Ser)_3_ linker, and expressed from the pZA31 plasmid (Figure SI 4A). We found that the addition of the (Gly-Ser)_3_ linker was critical to function, but we did not optimize the linker length.

**Figure 5.**
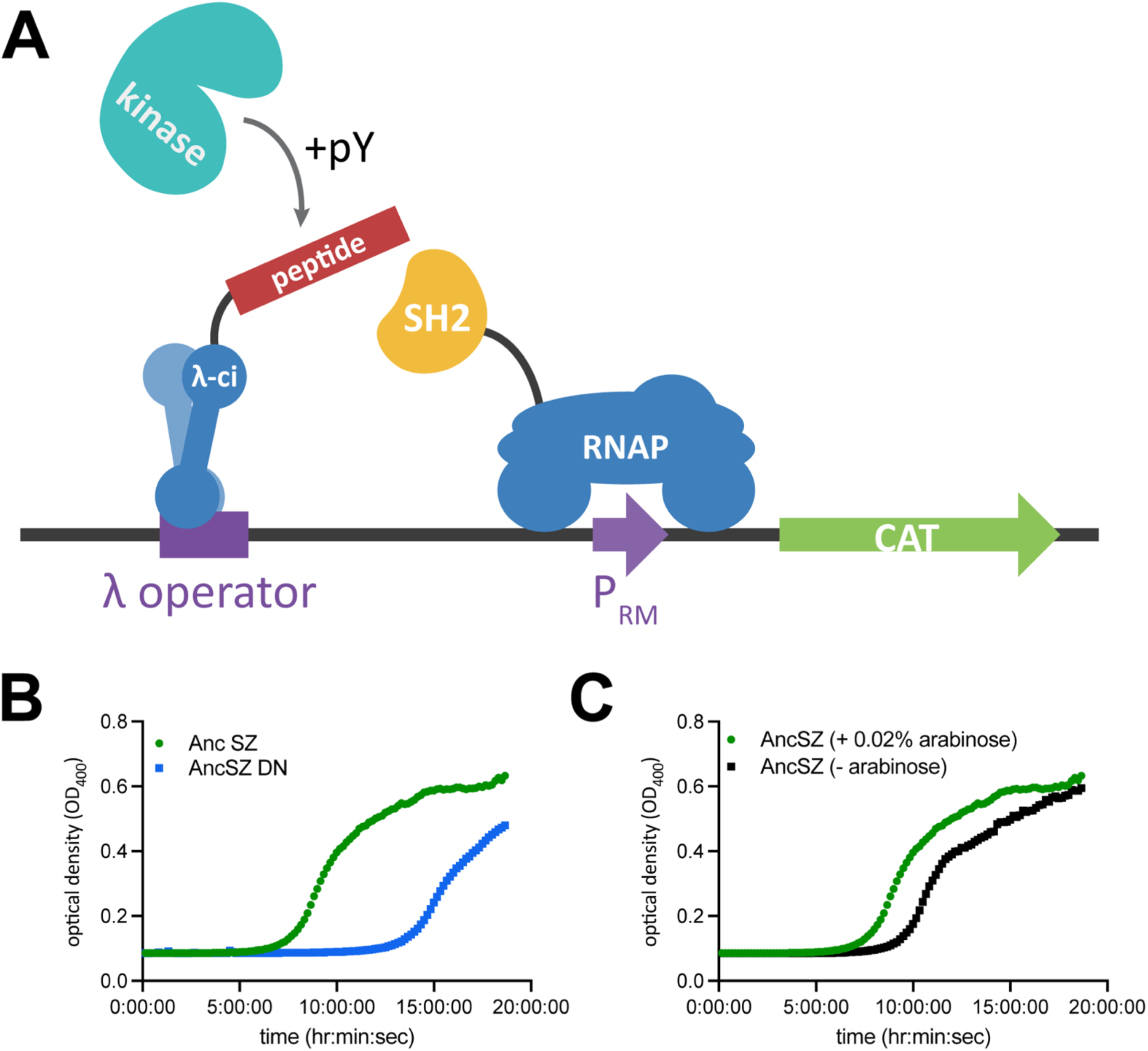
A bacterial two-hybrid for kinase activity. **A**. The protein tyrosine kinase phosphorylates a tyrosine on a peptide substrate, the “bait” protein, fused to the N-terminal domain of λ-cI which binds to an operator upstream of the phage λ-promoter (P_RM_). The “prey” protein, the Grb2 SH2 domain, binds the phosphorylated peptide, thereby recruiting the α-subunit of RNA polymerase to the promoter. Once at the promoter, RNA polymerase transcribes chloramphenicol acyl-transferase (CAT), which confers resistance to the antibiotic chloramphenicol. **B**. AncSZ begins growing well before the kinase dead mutant (AncSZ DN) in the presence of 50ng/µl chloramphenicol. **C**. Expression of the kinase is necessary for growth in chloramphenicol. When no arabinose is added the cells begin growing approximately 2 hours after those with 0.2% arabinose.

The reporter for kinase activity in this assay is the enzyme chloramphenicol acyltransferase (CAT), which confers resistance to the antibiotic chloramphenicol and is expressed from the phage λ-promoter (P_RM_). We opted to combine the kinase and reporter genes on one plasmid. The overexpression of active tyrosine kinases in *E. coli* is typically facilitated by the co-expression of a phosphatase, since protein tyrosine kinase activity can be toxic to bacteria.^6^ Instead of co-expressing a phosphatase, which would subvert the assay, we sought to maintain low levels of kinase expression, thereby potentially attenuating any cell toxicity caused by tyrosine kinase activity. To achieve this, we expressed the kinase using the titratable araBAD promoter, modified from the common expression vector pBAD.^42^ Figure SI 4B shows the map for this new plasmid, which will be referred to as pBpZR.

To determine whether bacterial growth in the presence of chloramphenicol was dependent on kinase activity, we initially tested two kinase variants, AncSZ and a mutant, AncSZ D486N, in which a catalytic aspartate residue, a conserved His-Arg-Asp (HRD) motif, was mutated to asparagine, rendering the kinase inactive. When the components of the bacterial two-hybrid were expressed in *E. coli*, the cultures transformed with the AncSZ kinase domain began growing in the presence of chloramphenicol earlier than those expressing the catalytically inactive kinase, indicating that bacterial growth was dependent on kinase activity (Figure 5B). The eventual growth of bacteria expressing the inactive kinase might be caused by recruitment of the RNA polymerase through non-specific interactions with the LAT peptide. Growth was dependent on arabinose; samples containing the kinase but not induced with arabinose showed delayed growth compared to the flasks containing 0.2% arabinose (Figure 5C).

### 2.4 Saturation mutagenesis of the AncSZ kinase domain

We generated a single-site saturation mutagenesis library of the AncSZ kinase domain. To achieve sufficient sequencing coverage of the entire kinase domain, we generated three sub-libraries (Pools 1-3 in Figure 6) in which 100 codon stretches of the kinase gene were mutagenized using degenerate primers to introduce NNS codons (N: A,C,T, or G and S: C or G) at each position in the kinase, theoretically resulting in 32 possible codons representing all 20 amino acids, including synonymous codons, and an amber stop codon. For each selection experiment, we transformed electrocompetent *E. coli* cells, which already contained the other two plasmids required for the two-hybrid assay, with one of the three sub-libraries. Transformed cells were grown without selection for three hours to allow for protein expression and then diluted into two separate growth flasks, one with the antibiotic chloramphenicol (selected population) and one without (unselected population). At the end of a seven-hour growth, the DNA from both flasks was harvested and deep sequenced (see Methods 4.7).

**Figure 6.**
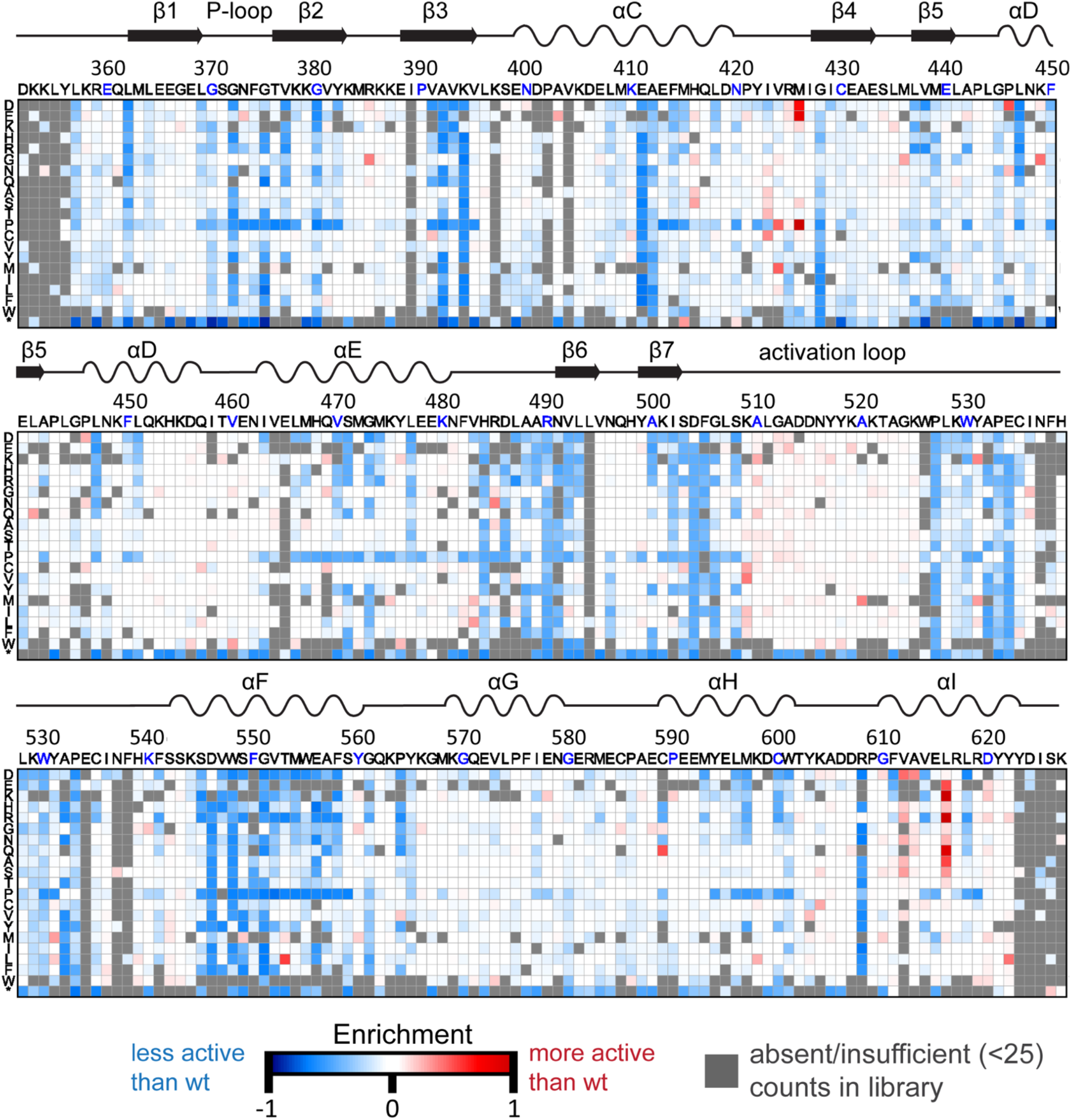
Saturation mutagenesis of AncSZ. Heatmap depicting the average enrichment values (E) for each pool (n=3). Along the top of each heatmap is the unmutated sequence of the protein, and along the left y-axis is the substituted residue. Synonymous codons are averaged. Substitutions which led to increased growth in chloramphenicol compared to the unmutated kinase are colored red. Those which resulted in decreased growth are blue. Grey boxes represent variants that were absent or had insufficient counts (<25) in the input library.

To quantify the relative activity of each variant in the library compared to the original AncSZ kinase, we calculated an enrichment score (Δ*E*_*sel,unsel*_):

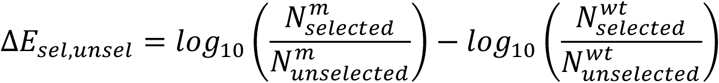

In this equation, 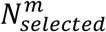 is the DNA count for that mutant in the selected population and 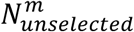 is the DNA count for the same mutant in the unselected population. The second term in the equation includes the “wild-type” counts, or the counts for the unmutated AncSZ sequence, in both the selected and unselected population. A negative value for Δ*E*_*sel,unsel*_ implies that the AncSZ variant of interest is less active than the original AncSZ, while a positive value suggests that it is more active. Because the activity of tyrosine kinases can be toxic to bacteria, we used the bacteria in the unselected flask as the reference sample. We envisioned that this would control for potential depletion of significant gain-of-function mutations due to cell death caused by the toxic effects of off-target kinase activity.

The Δ*E*_*sel,unsel*_ values for each kinase variant in the final library were calculated and plotted as a heatmap (Figure 5) using a custom Python script. All Δ*E*_*sel,unsel*_ values in Figure 5 represent the average across three replicates. Enrichment scores were reproducible across replicates (Figure SI 5). Modest differences between replicates can likely be attributed to the fact that the bacterial two-hybrid experiments are highly sensitive to antibiotic concentration, temperature, and inducer concentration. In the final libraries, not every amino acid substitution was observed or sufficiently abundant at each position, due to lack of amplification with the NNS primers or loss of this variant during one of library construction steps (see Methods 4.6). These absent variants are colored grey in Figure 6. Positive enrichment values, meaning that cells expressing the kinase variant grew better than those expressing original AncSZ, are red in Figure 5, while negative enrichment values are blue. Neutral variants, which the Δ*E*_*sel,unsel*_ values are close to zero, are colored white.

As seen in Figure 5, mutations at many positions are nearly neutral, supporting the principle that most proteins are robust to mutation. ^40,43-44^ An example of a residue that tolerates mutation is His104, an exposed surface residue (Figure SI 6A). Mutation of this residue to anything other than a stop codon is nearly neutral with the magnitude of Δ*E*_*sel,unse*_ within the noise of the experiment (Figure SI 6B).

### 2.5 Residues identified as loss-of-function are known to be critical for kinase function and/or structure

The site-saturation mutagenesis data identified many positions that are sensitive to mutation. In Figure 5, these positions are colored in shades of blue depending on the extent of the loss of function. Those positions that are the least tolerant of mutations, for which almost any substitution results in a loss-of-function (Figure 7A, right), overlap with many of the positions that are conserved across eukaryotic protein kinase families (Figure 7A, left).^1^ The locations of these mutations span both lobes of the catalytic domain, and many are critical for maintaining the active conformation of the kinase domain or are important for ATP binding.

**Figure 7.**
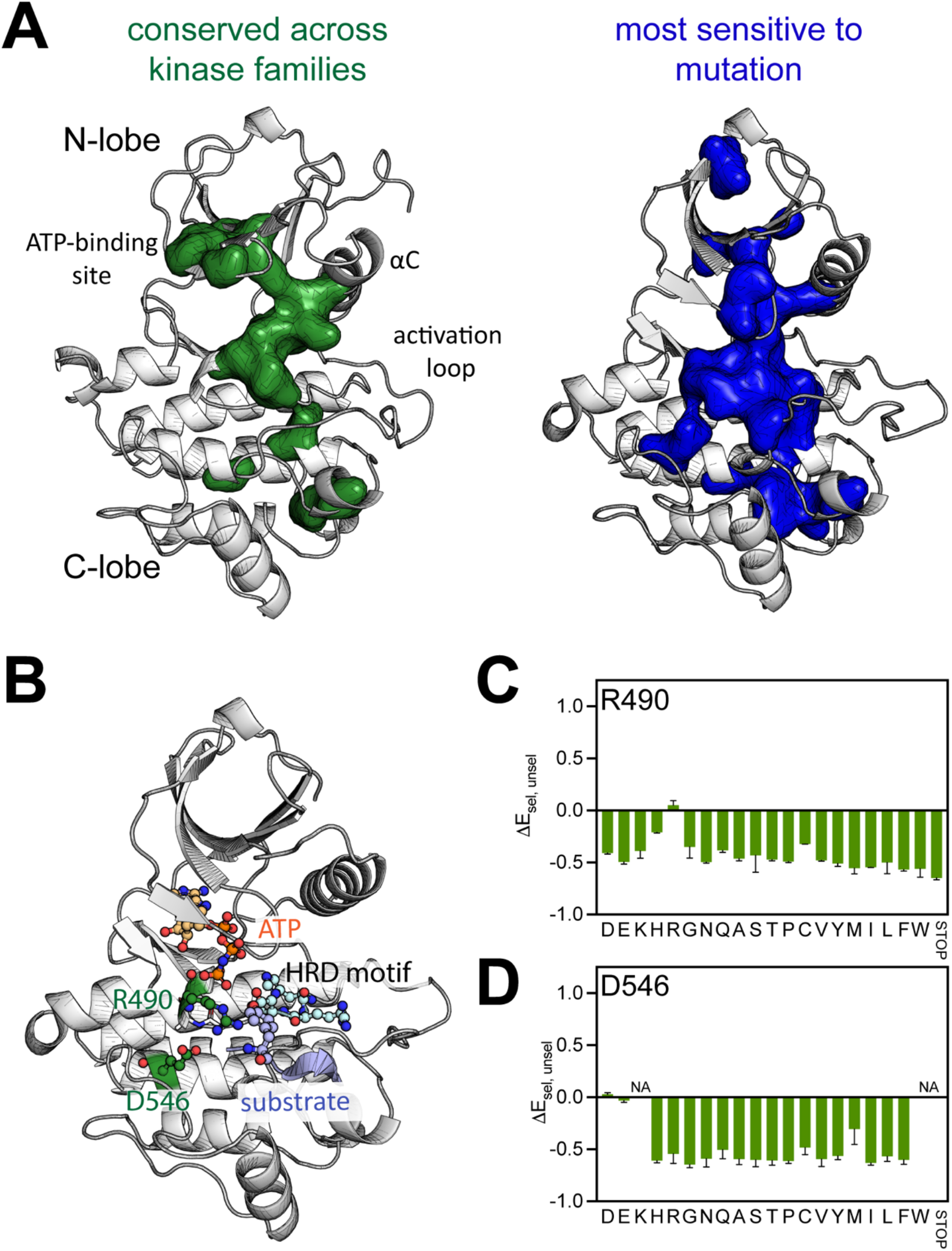
Loss-of-function mutations. **A**. Green residues (left) are invariant across all eukaryotic protein kinases, mapped onto the structure of Lck (PDB: 3LCK). These include some active site residues and residues involved in other essential motifs. The blue residues (right) are those to which almost any substitution results in a loss of activity in the bacterial two-hybrid assay of AncSZ. Residues are mapped onto a homology model of AncSZ. **B**. Global loss-of-function mutations Arg 490 and Asp 546 play important roles in the activity and/or structure of the kinase. **C**. Average enrichment scores for Arg 490 in the bacterial two-hybrid (n=3). For Arg 490, all mutations, except for synonymous R codons, are loss-of-function. **D**. Average enrichment scores for Asp 546. All mutations to D546, except for the synonymous aspartate or glutamate, are loss-of-function.

One mutationally-sensitive residue is an arginine in the catalytic loop of the kinase domain (Arg 490 in AncSZ; highlighted in green in Figure 7B). Mutation of this residue to anything other than a synonymous arginine results in a loss-of-function in the deep mutagenesis data (Figure 7C). The interactions facilitated by this arginine help to align the reactants appropriately for catalysis.

In Syk-family kinases this residue is located four residues after the catalytic aspartate (Asp 486 in AncSZ) in the conserved HRD motif. A structure of the insulin receptor tyrosine kinase (PDB: 1IR3) provides a reference for the active state of tyrosine kinases.^45^ In insulin receptor tyrosine kinase, Arg 1136 corresponds to Arg 490 in AncSZ, and it makes hydrogen bonds with the catalytic aspartate as well as the phenol oxygen on the substrate tyrosine.^45,46^

Some of the residues that show a strong loss-of-function phenotype likely play a critical role in maintaining the fold of the protein, rather than being directly involved in catalysis. Many large, buried hydrophobic residues fall into this category. Additionally, many positions in the kinase are selectively loss-of-function if mutated to proline (Figure SI 7). These residues are within secondary structure elements, and the introduction of proline is expected to disrupt the stability of these structures. Near the N-terminus of the F-helix is an acidic residue, Asp 546 (Figure 7B), that interacts with the backbone of the catalytic loop, acting to staple important residues, such as Arg 490 and those in the HRD motif to the F-helix. Mutations to Asp 546 are loss-of-function, except for glutamate or synonymous aspartate (Figure 7D).

### 2.6 Residues identified as gain-of-function are in regions known to regulate kinase activity

Many of the kinase variants identified as gain-of-function in the bacterial two-hybrid assay have mutations in regions known to be important for the allosteric regulation of kinase activity (Figure 8A). These regions include the activation loop (residues 504-530) and the αC-β4 loop (residues 418-427). We chose one of the variants in the αC-β4 loop, M426E, for further characterization (Figure 8B). We purified the M426E AncSZ kinase domain and tested its activity, both with and without pre-phosphorylation with Lck. Our data confirm that the purified AncSZ M426E has a higher kinase activity than AncSZ, by approximately 2.5-fold in the unphosphorylated state (Figure 7C).

**Figure 8.**
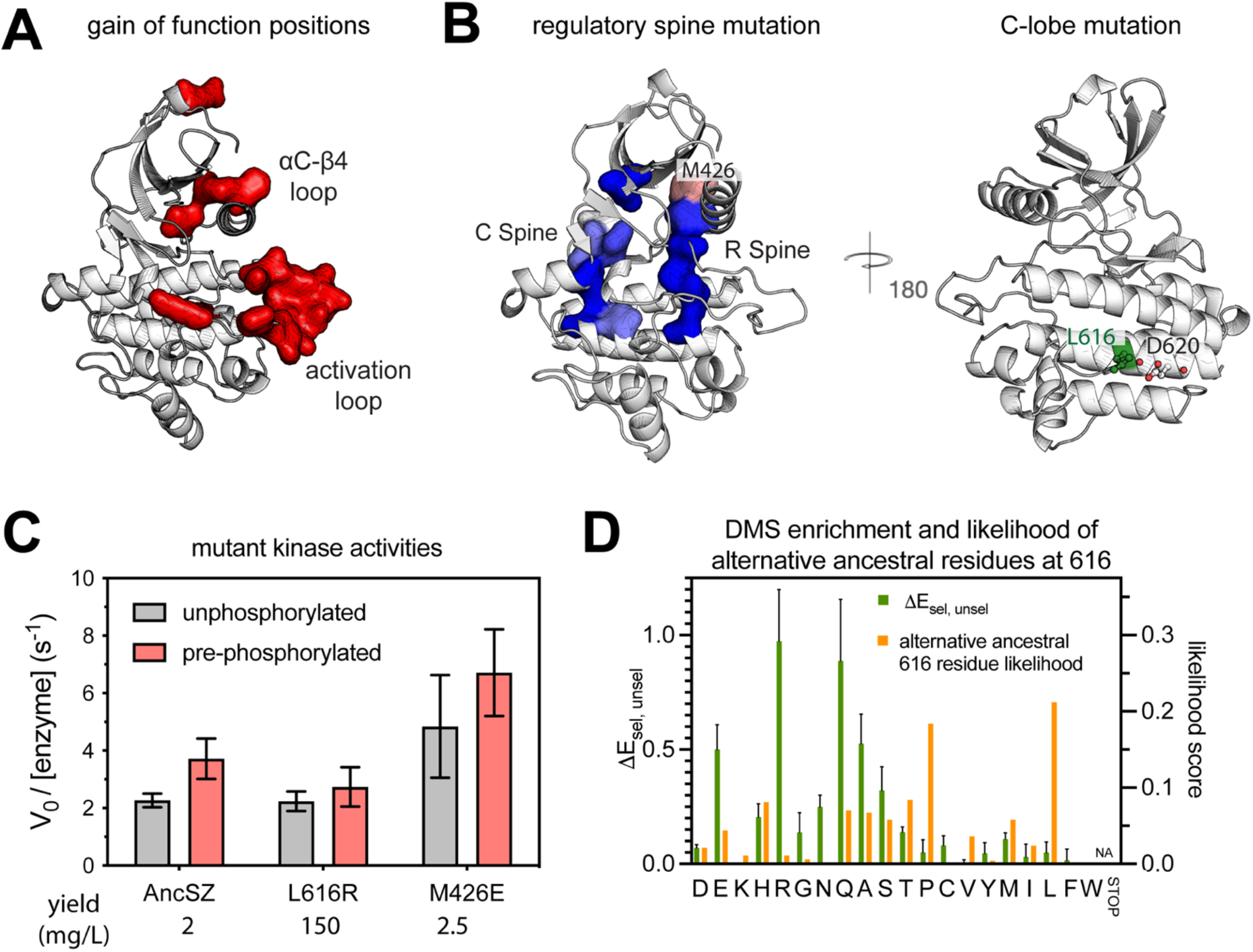
Activating mutations occur in regions important for activity but may also be the result of increased expression. **A**. Strong gain-of-function variants map to regions, such as the activation and αC-β4 loops, known to regulate kinase activity. **B**. Left, residues making up the catalytic spine (C-spine, left) and the regulatory spine (R-spine, right). Substitutions to these residues are loss-of-function in the bacterial two-hybrid assay, and thus colored blue. The top of the regulatory spine, Met 426, however, is a gain-of-function if mutated to Glu, Asp, or Pro and neutral for most other substitutions. Right, solvent-exposed Leu 616 is on the C-terminal helix of AncSZ. Next to L616 is D620 suggesting a salt bridge may form when the leucine is mutated to a positively charged arginine. **C**. AncSZ M426E has a higher LAT226 phosphorylation rate than the AncSZ or AncSZ* (AncSZ L616R). The red bars correspond to the rates determined following a one-hour treatment with a purified Lck kinase in order pre-phosphorylate the activation loop while the grey bars are the rates measured without this pre-phosphorylation. The yield observed during purification (n=1) for each kinase domain. The yield for AncSZ* was 75-fold higher than the other kinases. **D**. The average enrichment value for Leu 616 variants, left y-axis, and the likelihood score predicted by ancestral sequence reconstruction, right y-axis. Many of the variants that are gain-of-function in our assay were alternative amino acids at this position.

Met 426 is located at the top of the regulatory (R) spine, one of two hydrophobic spines that, along with the catalytic (C) spine, plays a critical role in the regulation of kinase activity.^47^ Assembly of these spines is a hallmark of the active conformation of protein tyrosine kinase domains.^47^ Overall, substitutions to these spines are broadly loss-of-function (Figure 8B), except for Met 426. Mutation of Met 426 to aspartate, glutamate, or proline results in a strong gain-of-function. These mutations are unlikely to stabilize the hydrophobic spine, and the mechanism by which they activate the kinase domain requires further study. In many tyrosine kinases the switch between an inactive and active conformation occurs through alterations in the orientation of helix αC. The αC-β4 loop modulates the conformation of helix αC, and in other kinases, such as EGFR, mutations to this loop underlie oncogenic activation.^5,48,49^ Met 426 is located at the end of this loop, near the beginning of β4.

### 2.7 Some gain-of-function mutants do not affect catalytic activity but instead enhance protein expression

One of the mutations in AncSZ that results in a large gain of function, L616R, is located on the face of the kinase domain opposite the active site, at a position that is not implicated in any catalytic or regulatory function. As mentioned above, during the expression and purification of the kinase domain of this variant, which was done with co-expression of the YopH phosphatase, we noted that the yield was ∼75-fold greater than for the original AncSZ (Figure 3A). The catalytic activity of this super-expressing variant (dubbed AncSZ*) is the same as that of the original AncSZ protein when measured *in vitro* using a purified LAT peptide (Figure 8C). The results of the screen show that mutation of this surface-exposed leucine in AncSZ to almost any of the hydrophilic residues results in gain-of-function (Figure 8D). These results reflect the interplay between stability, expression level, and fitness. The fusion of a fluorescent protein to AncSZ variants could allow for high-throughput monitoring of the expression of folded and soluble protein, as has been previously demonstrated.^50^

Ancestral sequence reconstruction does not predict a single ancestral sequence. Instead, it calculates the probability of an amino acid at each position. When generating the sequences for AncSZ, AncZ, and AncS, we chose the most probable amino acid at each position, based on the multiple sequence alignment and phylogenetic tree. However, previous work characterizing combinatorial libraries of the many putative ancestral states suggest that the alternative amino acids may affect the structure and/or function of the predicted protein.^51^ We re-visited the likelihoods generated by the ancestral sequence reconstruction for position 616 in AncSZ to determine whether any of the gain-of-function mutants identified were among the high probability amino acids in the sequence reconstruction. At position 616 the most probable amino acid was leucine, closely followed by proline. Other amino acids, including threonine, serine, glutamine, histidine, and alanine also had significant probabilities at this position (Figure 8D). While the most enriched variant, L616R, is not predicted at this position, many of those that are predicted also show up as gain-of-function in the bacterial two-hybrid assay (Figure 8D). These data support the need for testing multiple predicted ancestral sequences and suggest that the construction of ancestral libraries could further help in the identification of variants that are more readily expressed.

## 4. CONCLUSIONS

Several kinase families have been subject to ancestral sequence reconstruction, and the characterization of the predicted ancestral kinases has provided key insights into substrate specificity,^52^ regulation,^53^ and inhibitor binding.^54^ Here, we illustrate another use for ancestral sequence reconstruction in the study of protein kinases, which is the generation of variants with potentially improved yields when expressed in bacteria. The bacterially-expressed Syk-family kinase, AncSZ, retains general aspects of the catalytic activity, substrate specificity, and regulatory properties of the naturally occurring, extant Syk-family kinases. We developed a novel bacterial two-hybrid assay for Syk-family kinase activity and used it to carry out saturation mutagenesis of a Syk-family kinase. This deep mutational scan identified an apparent gain-of-function variant, AncSZ* (AncSZ L616R), which further biochemical characterization revealed is a super-expressing AncSZ variant with a comparable catalytical activity (Figure 8C) but even higher bacterial expression than the original AncSZ (Figure 3A).

The AncSZ kinase is sensitive to mutations at positions that are highly conserved in eukaryotic protein kinases. We identified gain-of-function mutations occurring in regions that are involved in regulating the transitions between active and inactive conformations, such as the αC-β4 loop. The identification of activating mutations implies that, even in the absence of other eukaryotic regulatory proteins and its own regulatory tandem SH2 module, which attenuate kinase activity *in vivo*, the isolated kinase domain is not maximally active. This may suggest that the conformational landscapes of Syk-family kinases have been tuned towards an inhibited state. Further biophysical and biochemical characterization of the ancestral proteins generated in this study, as well as activity-altering point mutations, may illuminate how this evolutionary tuning has been achieved and how it differs between Syk and ZAP-70.

The bacterial two-hybrid assay described here is easily generalizable, and, in principle, could be adapted to study any bacterially-expressed tyrosine kinase. It remains to be seen whether the low expression levels achieved with the titratable promoter will be successful in attenuating cell toxicity for kinases with less stringent substrate specificities than AncSZ. It will also be important to choose an appropriate peptide substrate to act as the bait and its corresponding SH2 domain to act as the prey. Some kinases, such as Abl, have substrate specificities that are distinct from those of ZAP-70.^36,55^ In the case of Lck, the Src family kinase upstream of ZAP-70 in T cells, the peptide substrate specificity is orthogonal to that of ZAP-70.^36^ Ideally it will be relatively easy to swap in the preferred substrate for the kinase being assayed, but some optimization will likely be necessary. Not only will this assay facilitate the construction and evaluation of many large libraries of kinase variants, but it may also enable the identification of new soluble versions of difficult to express kinases. These soluble, model kinases would be powerful tools for the high-throughput study of the structure, function, and inhibition of protein tyrosine kinases.

## 4. METHODS AND MATERIALS

### 4.1 Ancestral Sequence Reconstruction

The ancestral sequence reconstruction to generate the AncSZ, AncS, and AncZ kinase sequences was carried out in three steps: (1) construction of a multiple sequence alignment, (2) construction of a phylogenetic tree, and (3) prediction of ancestral states. The multiple sequence alignment was created as described previously.^**27**,**33**^ Briefly, the full-length human Syk and ZAP-70 sequences were used as query sequences in a series BLAST searches within annotated metazoan proteomes.^**56**^ To extend our search, once distantly related sequences (e.g. those from fish species) were identified, those were used as queries for additional searches. As noted previously, most organisms contained two Syk-family kinases, which could be reliably designated as Syk or ZAP-70 based on higher homology to one human kinase or the other.^**27**^ For invertebrates and jawless vertebrates, only a single Syk-family kinase was typically identified. All of the sequences that we compiled had the expected tandem-SH2/kinase domain architecture, and any partial sequences were omitted from downstream analyses. In total, we had 89 Syk orthologs and 87 ZAP-70 orthologs from jawed vertebrates, and 7 sequences from invertebrates and jawless vertebrates (183 sequences in total).

A multiple sequence alignment was generated using the software T-Coffee, followed by small manual adjustments in linker regions and at the C-terminus. The multiple sequence alignment is provided as Supplementary File 1. The multiple sequence alignment was then used to construct a phylogenetic tree using the PHYLIP package,^**57**^ using the Syk-family kinase sequence from *Amphimedon queenslandica* as the outgroup. In this tree, the jawed-vertebrate Syk and ZAP-70 sequences clustered into two distinct clades that had internal structures consistent with known species relationships. This phylogenetic tree is provided as Supplementary File 2. Finally, the multiple sequence alignment and phylogenetic tree were used as input for the software Lazarus.^**30**^ In order to assign the sequences of the predicted ancestors of the Syk and ZAP-70 lineages, as well as their common ancestor (AncS, AncZ, and AncSZ, respectively), we identified the relevant internal nodes and selected the most like amino acid at each position.

### 4.2 Protein Constructs and Purification

Ancestral kinase genes designed from the predicted protein sequence, codon-optimized for expression in *E. coli*. The sequences of the genes are attached as Supplementary File 3. The genes were synthesized by IDT. The isolated kinase domains genes were cloned with an uncleavable C-terminal His_6_ tag, and the full-length kinases were cloned with a C-terminal PreScission protease-cleavable His_6_ tag. ZAP-70, Syk, AncZ, and AncS constructs were all cloned into the pFastBac1 vector, and the resulting plasmids were used to generate bacmids in DH10Bac cells. Bacmids were then used to transfect Sf21 insect cells using standard protocols. Baculovirus harvested from the cell culture supernatant was used to infect 4-6 L of Sf21 cells, and protein expression was carried out for 2-4 days. The AncSZ constructs were cloned into the pET-23a vector. The AncSZ proteins were co-expressed at 18 °C in 2L of BL21(DE3) cells with the tyrosine phosphatase YopH.

After overexpression, insect or bacterial cells were suspended in Tris buffer (50 mM, pH 8.0) containing 300 mM NaCl, 10 mM imidazole, 2 mM 2-mercaptoethanol, 10% glycerol, and a cocktail of protease inhibitors. Cells were lysed using a cell homogenizer, and lysates were clarified by ultracentrifugation. The supernatant was filtered then purified over nickel and ion exchange columns. At this stage, the proteins that were expressed in insect cells were treated with purified YopH to ensure they were fully dephosphorylated, whereas those co-expressed with YopH in *E. coli* were isolated in their dephosphorylated state. Full-length proteins were treated with PreScission protease to remove the C-terminal His_6_ tags. Finally, all of the proteins were purified by size exclusion chromatography into their storage buffer: 10 mM HEPES, pH 7.5, 150 mM NaCl, 5 mM MgCl_2_, 1 mM TCEP, and 10% glycerol. Purified proteins were flash frozen in small aliquots at concentrations ranging from 20-400 μM and stored at -80 °C.

### 4.3 In vitro kinase activity assays

*In vitro* peptide phosphorylation measurements were carried out using a continuous colorimetric assay, as described previously.^**33**^ The production and purification of all peptides used for these measurements was also described previously.^**33**^ For all measurements with the isolated kinase domain, the peptide concentration was 500 μM, but kinase concentration was varied as follows: 1 μM for ZAP-70 and AncZ, 0.2 μM for AncSZ, AncS, and Syk. In order to measure the effects of pre-phosphorylation, each Syk-family kinase, at a concentration of 7.5 μM, was treated with 1 mM ATP and 0.75 μM Lck kinase domain for one hour. Then, the pre-phosphorylation mixture was diluted to concentrations described above to measure phosphorylation of the LAT Y226 peptide. The Lck kinase domain, on its own, showed a negligible rate of LAT peptide phosphorylation under the same conditions, confirming that all of the measured activity in these experiments came from Syk-family kinases.

For auto-activation assays with full-length Syk, ZAP-70, and AncSZ, the kinases (500 nM) were incubated with the LAT Y226 peptide (250 μM), and ADP production was monitored with the same continuous colorimetric assay described above. As a reference for the fully activated states of the kinases, pre-activated samples were prepared by treating the Syk-family kinases (3.75 μM) with 1 mM ATP and the Lck kinase domain (0.375 μM) for 1 hour. After Lck-mediated activation, the pre-activation reactions were diluted into the LAT Y226 phosphorylation reaction mixtures and analyzed alongside un-activated Syk-family kinases.

The activation loop phosphorylation states of all kinases used for *in vitro* assays were assessed by western blotting. Proteins were transferred to PVDF membranes using a semi-dry transfer apparatus with CAPS transfer buffer (10 mM CAPS, pH 11, 10% MeOH). Membranes were probed with Cell Signaling Technologies Phospho-Zap-70 (Tyr493)/Syk (Tyr526) Antibody #2704 at a dilution of 1:2000, followed by Cell Signaling Technologies anti-rabbit IgG HRP-conjugated secondary antibody at a dilution of 1:1000. Blots treated with enhanced chemiluminescence reagents and imaged. We note that the phospho-specific antibody for the Syk/ZAP-70 activation loops shows off-target reactivity toward Lck, but the Lck construct could be readily distinguished from Syk-family kinases based on its size. For autophosphorylation experiments, the kinases were mixed with 1 mM ATP at 500 nM or 5 μM. At various time points, the reactions were quenched with SDS-PAGE loading dye containing 5 mM EDTA. Samples were run on gel. Western blots were conducted as described above.

### 4.4. Substrate specificity screen

All screens were carried out as described in Shah et al. ^**33**^ In all cases, the isolated kinase domains were used at a concentration of 500 nM. To achieve similar library phosphorylation levels across the kinases, library phosphorylation reactions were carried out for different amounts of time, as follows: ZAP-70 for 30 minutes, AncZ for 15 minutes, AncSZ for 3 minutes, AncS for 3 minutes, and Syk for 3 minutes.

### 4.5 Construction of pBpZR Plasmid

The pBAD LIC cloning vector (8A) was a gift from Scott Gradia (Addgene plasmid # 37501 ; http://n2t.net/addgene:37501 ; RRID:Addgene_37501).

The following primers were used for the Gibson Assembly of the pBAD and pZERM plasmid components:

pZERM_Fwd-aacattgaaaaaggaagagtcagctcactcaaaggcggtaatacggttatccac

pZERM_Rev-gacgcatcgtggccggcatccggccgcttacgccccgccc

pBAD_Fwd-gggcggggcgtaagcggccggatgccggccacgatgcgtc

pBAD_Rev-taccgcctttgagtgagctgactcttcctttttcaatgttattgaagcatttatcag

The RBS and restriction enzyme sites were introduced via the following primer pair and assembled using Golden Gate assembly:

AncSZ_rbs_fwd-ggaggagggtctcactaatttgtttaactttaagaaggagacatctagaatggacaagaaactttacctgaaacgc

AncSZ_rbs_rev-ggaggagggtctcattagcccaaaaaaacgggtatggagaaacag

### 4.6 Construction of Saturation Mutagenesis Library

The saturation mutagenesis library of AncSZ was constructed using oligonucleotide-directed mutagenesis of the gene encoded on the pET27B plasmid. For each amino acid position two primers were generated, a sense and anti-sense, which when used together amplify the entire plasmid as well as introduce two BsaI restriction enzyme sites thereby enabling Golden Gate Assembly of the mutated plasmid. The anti-sense primer of each pair contained the degenerate NNS codon, where N is a mixture of A, C, G and T nucleotides and S is a mix of C and G nucleotides. This degenerate codon allows for one primer pair to introduce up to 32 possible codons at each position. These 32 codons comprise all 20 amino acids as well as a stop codon. Each primer pair was used in separate PCRs. Following amplification each reaction was gel verified. Successful products were pooled in three libraries (Pool 1: Residues 2-100, Pool 2: Residues 90-189, Pool 3: Residues 178-278) for subsequent gel purification and Golden Gate assembly with BsaI (NEB) and T4 DNA ligase (NEB).

NEB® 10-beta electrocompetent *E. coli* were transformed with each library to get >100X coverage for each pool. Successful pools were miniprepped to obtain the library in pET27B. This library was then digested with XbaI and BamHI-HF restriction enzymes. The pBpZR plasmid was similarly digested. Both were gel purified. The pBpZR plasmid and digested kinase library were ligated using T4 DNA ligase overnight at 4C. The ligated product was transformed into NEB® 10-beta electrocompetent *E. coli* to get >100X coverage for each pool. If the coverage was sufficient the cells were miniprepped to obtain the library in the pBpZR expression vector. Some mutations did not make it through this pipeline and therefore are not represented in the final data set.

### 4.7 Bacterial Two-Hybrid Assay

The library was transformed into electrocompetent Bl21.DE3 cells already containing the Grb2 SH2-RNAP and Lambda CI-LAT226 constructs, ensuring 100X coverage. The assay was originally developed using MC4100-Z1 cells, which have a z1 cassette expressing the repressors LacI and TetR, but for this study the assay was carried out in BL21.DE3 cells.^25,40^ We did not change induction protocol when working with BL21.DE3 cells. The use of the original MC4100-Z1 cells and optimization of induction conditions could further improve the assay. A saturated overnight culture of transformed cells was diluted to an OD600 of 0.001 in media containing 20µg/mL trimethoprim, 50µg/mL kanamycin, 100µg/mL ampicillin, 50ng/mL doxycycline, and 0.02% arabinose. These induced cells were grown for three hours, at which point the OD600 was typically close to 0.1, to allow for expression of the necessary bacterial two-hybrid components. Following this three-hour induction period, the cells were again diluted to an OD600 of 0.001 into two flasks, the selected flask containing 50µg/mL chloramphenicol + antibiotics + inducers and the unselected flask containing just the antibiotics + inducers. Leftover cells from the initial three-hour growth were miniprepped in order to allow for sequencing of the population just prior to selection (T0). The cells were grown for an additional 7 hours. The unselected population typically was close to saturation following this growth while the selected population typically reached a final OD of just above 0.10. Both populations were miniprepped at the end of the growth period.

The selected and unselected DNA samples were prepared for sequencing through two, sequential PCR steps, each of which contains only 20 cycles of amplification to reduce any PCR introduced errors. First, ∼20ng of each DNA sample was used in a PCR using primers which amplified on the part of the gene which had been mutagenized (Pool 1, Pool 2, or Pool 3). These primers also included 5’ overhangs overlapping with Illumina adapter sequences to act as PCR handles in the subsequent amplification. In the second PCR, primers were used to introduce unique TruSeq indices for each sample and generate a ∼450bp amplicon for sequencing on the MiSeq sequencer using a 500 cycles kit. The concentration of the PCR product was determined using Picogreen (Thermofisher) and denatured prior to sequencing. The final concentration loaded onto the MiSeq chip was 10pM of the pooled denatured DNA.

## ACKNOWLEDGEMENTS

This paper is dedicated to the memory of Danny Tawfik. One of us (JK) benefited enormously from a long conversation with Danny on approaches to high-throughput mutagenesis and *in vitro* evolution, prior to committing the lab to an increased use of these methods. We thank the members of the Marqusee and Kuriyan labs for helpful discussions. We are especially grateful to Subu Subramanian for his advice when constructing NNS libraries. This work was supported by the US NIH R01GM050945 to SM, the US NIH PO1 5P01AI091580-10 to JK and the Damon Runyon Cancer Research Foundation DRG-14 to NHS. JFA was supported by a Jane Coffins Childs Memorial Fund for Medical Research fellowship. SM is a Chan Zuckerberg Biohub Investigator. *This article is subject to HHMI’s Open Access to Publications policy. HHMI lab heads have previously granted a nonexclusive CC BY 4*.*0 license to the public and a sublicensable license to HHMI in their research articles. Pursuant to those licenses, the author-accepted manuscript of this article can be made freely available under a CC BY 4*.*0 license immediately upon publication*.

## Notes

### Competing Interest Statement

The authors have declared no competing interest.

