## Supplementary File 3 for "Saturation mutagenesis of a predicted ancestral Syk-family kinase"

**ZAP-70 near-full-length (residues 1-606)**

MPDPAAHLPFFYGSISRAEAEEHLKLAGMADGLFLLRQCLRSLGGYVLSLVHDVRFHHFPIERQLNGTYAIAGGKAHCGPAELCEFYSRDPDGLPCNLRKPCNRPSGLEPQPGVFDCLRDAMVRDYVRQTWKLEGEALEQAIISQAPQVEKLIATTAHERMPWYHSSLTREEAERKLYSGAQTDGKFLLRPRKEQGTYALSLIYGKTVYHYLISQDKAGKYCIPEGTKFDTLWQLVEYLKLKADGLIYCLKEACPNSSASNASGAAAPTLPAHPSTLTHPQRRIDTLNSDGYTPEPARITSPDKPRPMPMDTSVYESPYSDPEELKDKKLFLKRDNLLIADIELGCGNFGSVRQGVYRMRKKQIDVAIKVLKQGTEKADTEEMMREAQIMHQLDNPYIVRLIGVCQAEALMLVMEMAGGGPLHKFLVGKREEIPVSNVAELLHQVSMGMKYLEEKNFVHRDLAARNVLLVNRHYAKISDFGLSKALGADDSYYTARSAGKWPLKWYAPECINFRKFSSRSDVWSYGVTMWEALSYGQKPYKKMKGPEVMAFIEQGKRMECPPECPPELYALMSDCWIYKWEDRPDFLTVEQRMRACYYSLASKVEGGSGLEVLFQ|GPHHHHHH

**AncSZ full-length (residues 1-627)**

MADSANHLPYFYGSITREEAEDYLKQGGMSDGLFLLRQSLNSLGGYVLSVVYDRQCHHYTIERQLNGTYAIAGGKPHSGPAELCEYHSQDSDGLVCLLKKPCNRPPGVQPKVGPFEDLKDQLIREYVRQTWNLEGEALEQAIISQRPQLEKLIATTAHEKMPWFHGKISREESERRLLSGAQPNGKFLIRERDENGSYALSLLYEKKVYHYRIDRDKSGKLSIPDGKKFDTLWQLVEHYSHKPDGLLCVLTEPCPNPDSPAGALGAPAPPLPGSHPKLETAGGIISRIKSYSFPKPGFKKKPPSERPKSALNVNGYVPRPKPLGAEGGSRRAMPMDTNVYESPYSDPEELKDKKLYLKREQLMLEEGELGSGNFGTVKKGVYKMRKKEIPVAVKVLKSENDPAVKDELMKEAEFMHQLDNPYIVRMIGICEAESLMLVMELAPLGPLNKFLQKHKDQITVENIVELMHQVSMGMKYLEEKNFVHRDLAARNVLLVNQHYAKISDFGLSKALGADDNYYKAKTAGKWPLKWYAPECINFHKFSSKSDVWSFGVTMWEAFSYGQKPYKGMKGQEVLPFIENGERMECPAECPEEMYELMKDCWTYKADDRPGFVAVELRLRDYYYDISKGSGLEVLFQ|GPHHHHHH

**Syk full-length (residues 1-635, A2G mutation)**

MGSSGMADSANHLPFFFGNITREEAEDYLVQGGMSDGLYLLRQSRNYLGGFALSVAHGRKAHHYTIERELNGTYAIAGGRTHASPADLCHYHSQESDGLVCLLKKPFNRPQGVQPKTGPFEDLKENLIREYVKQTWNLQGQALEQAIISQKPQLEKLIATTAHEKMPWFHGKISREESEQIVLIGSKTNGKFLIRARDNNGSYALCLLHEGKVLHYRIDKDKTGKLSIPEGKKFDTLWQLVEHYSYKADGLLRVLTVPCQKIGTQGNVNFGGRPQLPGSHPATWSAGGIISRIKSYSFPKPGHRKSSPAQGNRQESTVSFNPYEPELAPWAADKGPQREALPMDTEVYESPYADPEEIRPKEVYLDRKLLTLEDKELGSGNFGTVKKGYYQMKKVVKTVAVKILKNEANDPALKDELLAEANVMQQLDNPYIVRMIGICEAESWMLVMEMAELGPLNKYLQQNRHVKDKNIIELVHQVSMGMKYLEESNFVHRDLAARNVLLVTQHYAKISDFGLSKALRADENYYKAQTHGKWPVKWYAPECINYYKFSSKSDVWSFGVLMWEAFSYGQKPYRGMKGSEVTAMLEKGERMGCPAGCPREMYDLMNLCWTYDVENRPGFAAVELRLRNYYYDVVNGSGLEVLFQ|GPHHHHHH

**ZAP-70 kinase domain (Met, residues 327-606)**

MDKKLFLKRDNLLIADIELGCGNFGSVRQGVYRMRKKQIDVAIKVLKQGTEKADTEEMMREAQIMHQLDNPYIVRLIGVCQAEALMLVMEMAGGGPLHKFLVGKREEIPVSNVAELLHQVSMGMKYLEEKNFVHRDLAARNVLLVNRHYAKISDFGLSKALGADDSYYTARSAGKWPLKWYAPECINFRKFSSRSDVWSYGVTMWEALSYGQKPYKKMKGPEVMAFIEQGKRMECPPECPPELYALMSDCWIYKWEDRPDFLTVEQRMRACYYSLASKVEGSHHHHHHG

**AncZ kinase domain (Met, residues 321-596)**

MDKKLFLKRDQLMIDEVELGSGNFGCVKKGVYKMRKKQIDVAIKVLKSENEKAVKDEMMKEAEIMHQLDNPYIVRMIGICQAESLMLVMEMAPGGPLNKFLSSKKDQITVENIVELMHQVSMGMKYLEEKNFVHRDLAARNVLLVNQHYAKISDFGLSKALGADDNYYKARTAGKWPLKWYAPECINFHKFSSKSDVWSFGVTMWEAFTYGQKPYKKMKGPEVISFIENGNRMECPADCPEEMYTLMKDCWTYKHEDRPGFVAVEERMRNYYYSISKHHHHHH

**AncSZ kinase domain (Met, residues 352-627)**

MDKKLYLKREQLMLEEGELGSGNFGTVKKGVYKMRKKEIPVAVKVLKSENDPAVKDELMKEAEFMHQLDNPYIVRMIGICEAESLMLVMELAPLGPLNKFLQKHKDQITVENIVELMHQVSMGMKYLEEKNFVHRDLAARNVLLVNQHYAKISDFGLSKALGADDNYYKAKTAGKWPLKWYAPECINFHKFSSKSDVWSFGVTMWEAFSYGQKPYKGMKGQEVLPFIENGERMECPAECPEEMYELMKDCWTYKADDRPGFVAVELRLRDYYYDISKHHHHHH

**AncS kinase domain (Met, residues 353-628)**

MEKTLYLKREQLILEEGELGSGNFGTVKKGIYKMRKKEIPVAVKVLKSENDPAVKDELMKEANFMHQLDNPYIVRMIGICEAESLMLVMELAELGPLNKFLQKHKDHITVKNIVELVHQVSMGMKYLEEKNFVHRDLAARNVLLVTQHYAKISDFGLSKALNADENYYKAKTTGKWPLKWYAPECINFFKFSSKSDVWSFGVMMWEAFSYGQKPYKGMKGQEVLPMIENGERMECPAKCPPEMYDLMKDCWTYKADDRPGFVAVEPRLRDYYYDISKHHHHHH

**Syk kinase domain (Met, residues 356-635)**

MEEIRPKEVYLDRKLLTLEDKELGSGNFGTVKKGYYQMKKVVKTVAVKILKNEANDPALKDELLAEANVMQQLDNPYIVRMIGICEAESWMLVMEMAELGPLNKYLQQNRHVKDKNIIELVHQVSMGMKYLEESNFVHRDLAARNVLLVTQHYAKISDFGLSKALRADENYYKAQTHGKWPVKWYAPECINYYKFSSKSDVWSFGVLMWEAFSYGQKPYRGMKGSEVTAMLEKGERMGCPAGCPREMYDLMNLCWTYDVENRPGFAAVELRLRNYYYDVVNSHHHHHHG
