## Supplementary Figures 1-7 for "Saturation mutagenesis of a predicted ancestral Syk-family kinase"

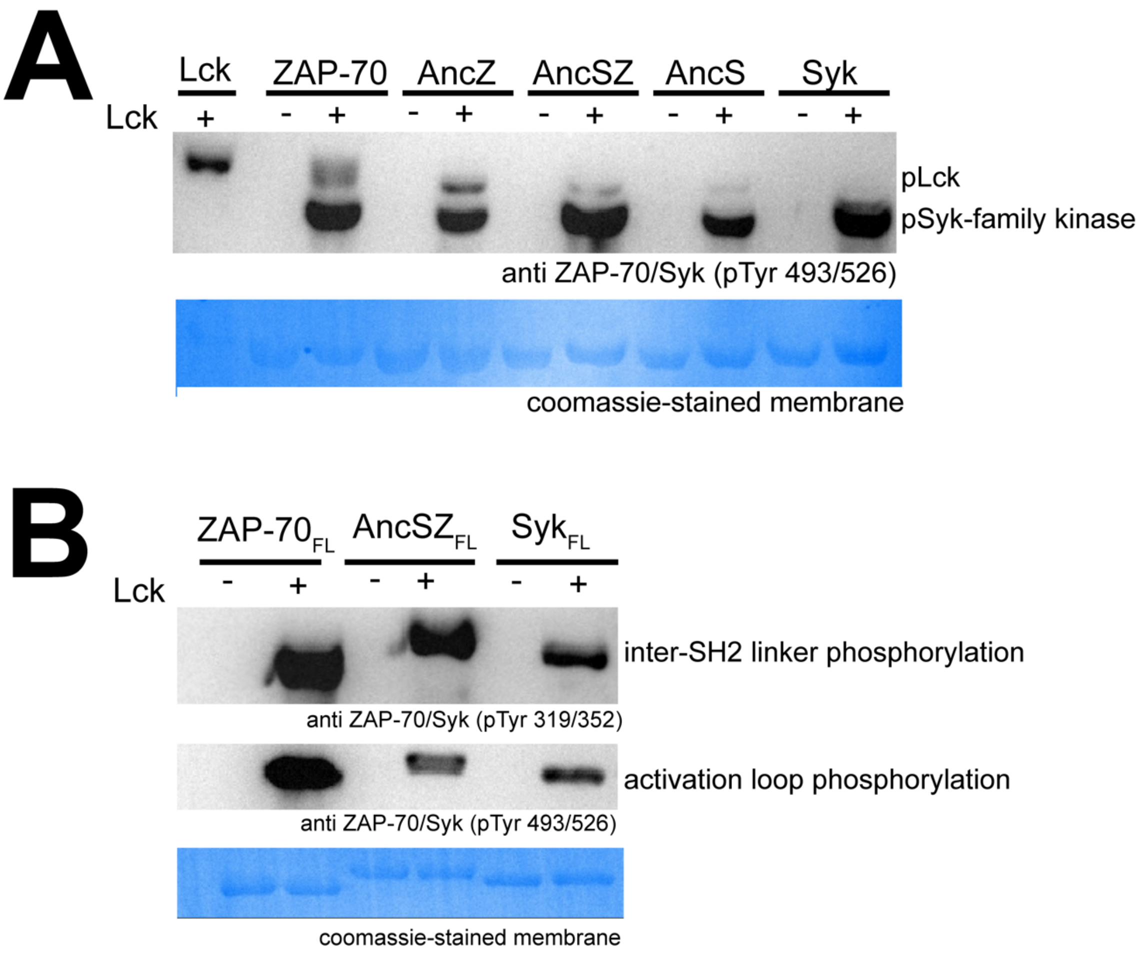


**Figure SI 1.** *Pre-phosphorylation of Syk-family kinases with Lck* **A.** Western blot of the five Syk-family kinases with and without incubation with the kinase Lck using a primary antibody that recognizes the phosphorylated activation loop of Syk-family kinases. The membranes were stained with Coomassie following imaging. **B.** Western blot of full-length Syk, ZAP-70, and AncSZ with and without incubation with Lck using primary antibodies recognizing the phosphorylated activation loop (bottom blot) and phosphorylated inter-SH2 linker (top blot). The membranes were stained with Coomassie following imaging.

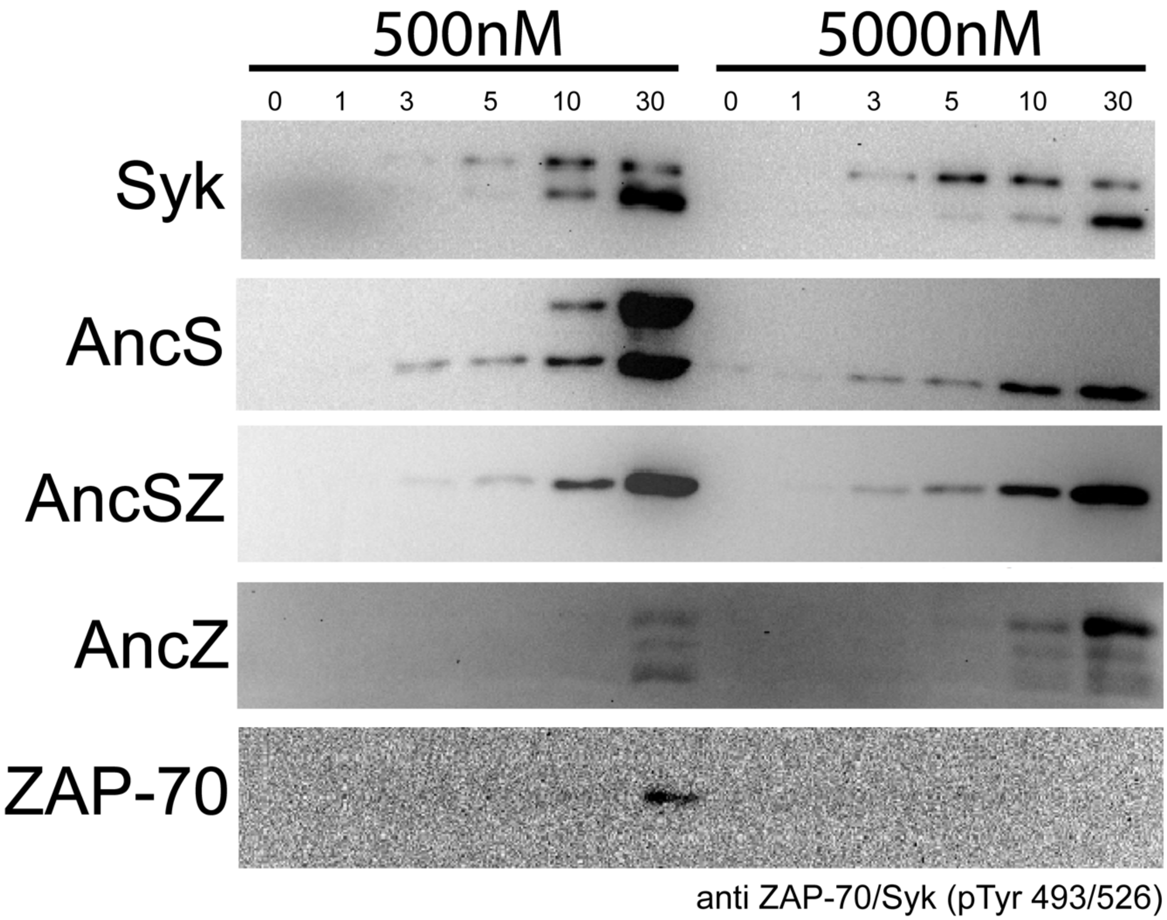


**Figure SI 2.** *Activation loop auto-phosphorylation by Syk-family kinases.* Western blots depicting the time course of auto-phosphorylation of the kinase domains of human Syk, AncS, AncSZ, AncZ, and human ZAP-70 using a primary antibody that recognizes the phosphorylated tyrosine on the activation loop(s). Auto-phosphorylation is slow for all the kinases, but especially so for AncZ and human ZAP-70.

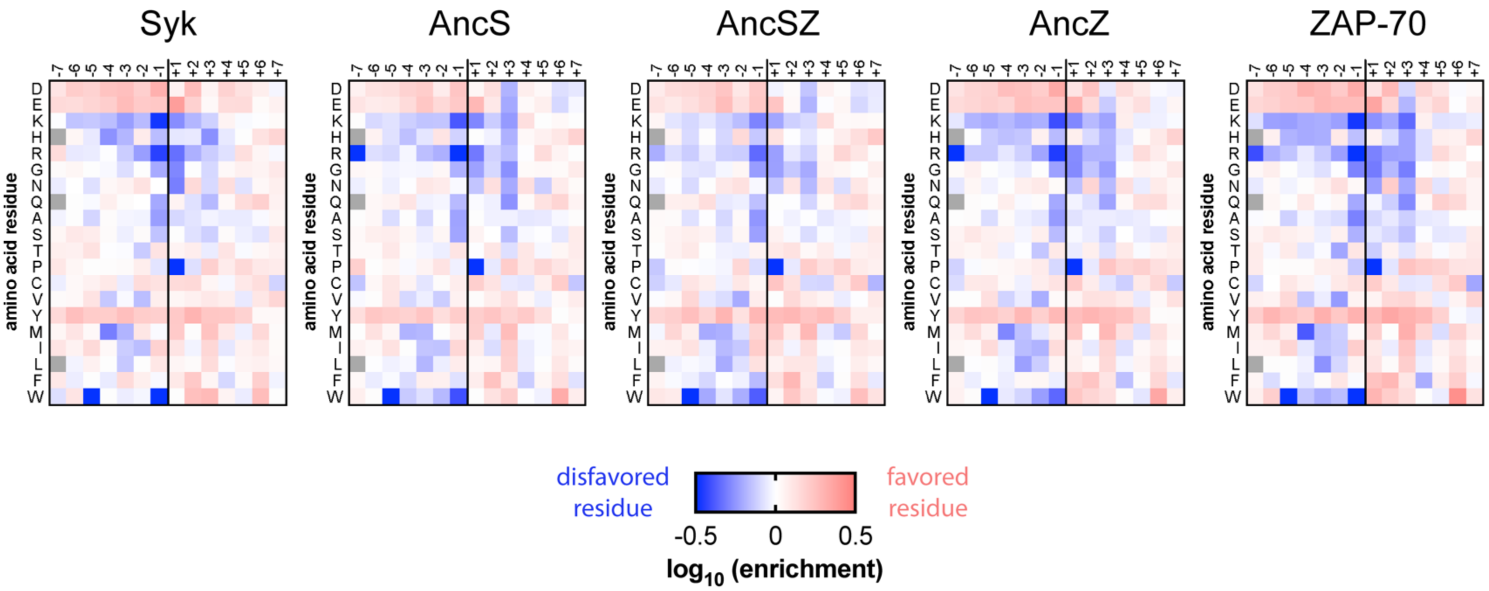


**Figure SI 3.** *The substrate specificity of human and ancestral Syk family kinases.* Heatmaps depicting the enrichment of amino acids at all positions in the substrate peptide for each Syk-family kinase, expanding on the data in Figure 4.

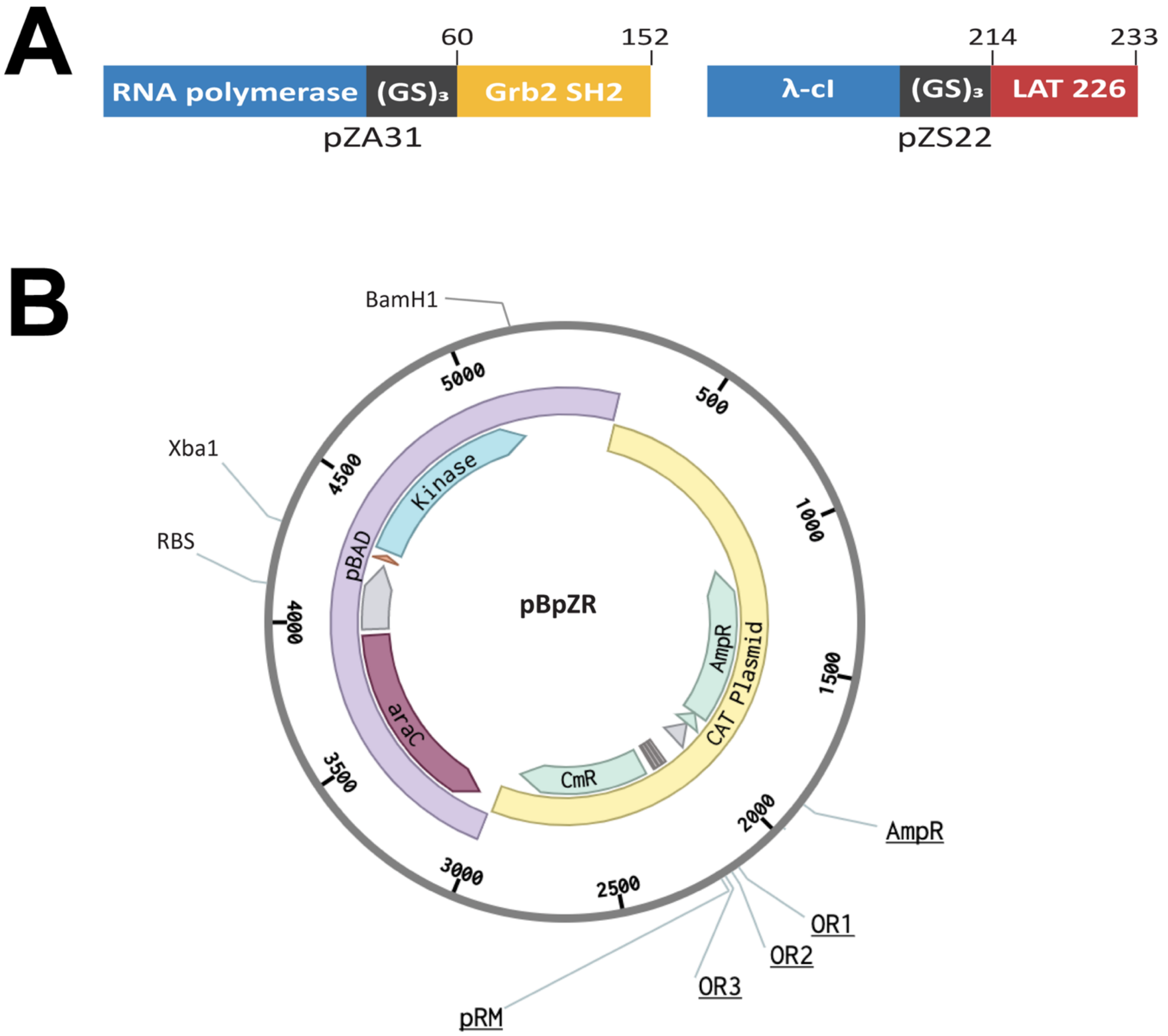


**Figure SI 4.** *Constructs used in bacterial two-hybrid* **A.** In the "prey" construct, RNA polymerase is followed by a Gly-Ser linker and then the Grb2 SH2 domain. In the "bait" construct, the N-terminal domain of the λ-cI protein is fused to the LAT 226 peptide by a Gly-Ser linker. Specific residue numbers used are denoted, and the expression vector is listed beneath each construct. **B.** The pBpZR vector that was constructed for the two-hybrid assay. The portion originating from the pBAD expression vector is shown in light purple, and the portion originating from the original CAT plasmids shown in yellow. The pBAD portion contains the araBAD promoter (grey), the corresponding araC gene, a ribosome binding site (orange), and inserted restriction enzyme sites (Xba1 and BamH1). The CAT portion contains the CAT gene, the phage promoter (PRM) and operons (OR1-3), and the beta-lactamase gene.

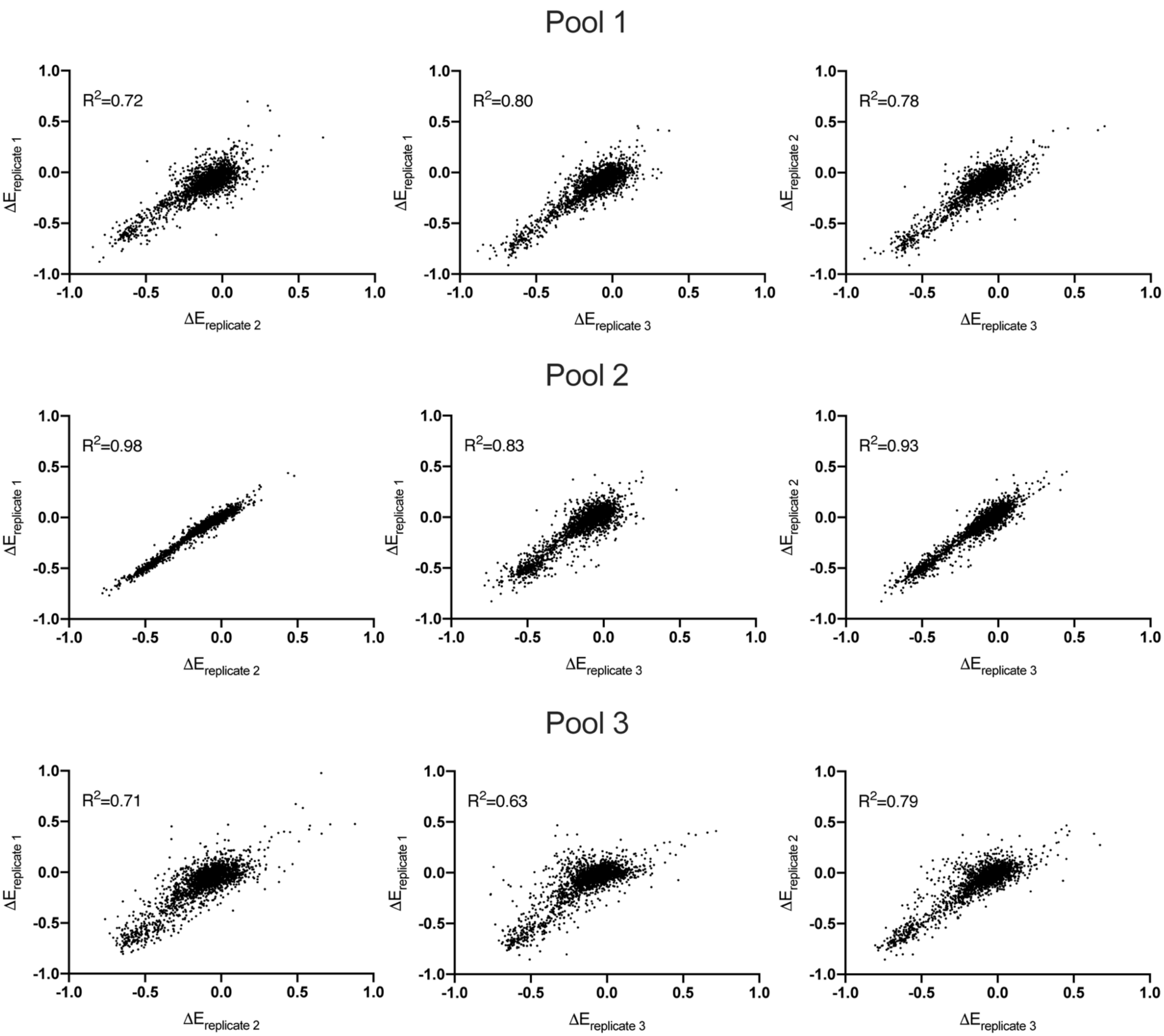


**Figure SI 5.** *The bacterial two-hybrid assay is reproducible.* The correlation of the enrichment scores for the selected population over the unselected populated for three independent replicates for each pool. All reported enrichment scores will be the average from three replicates.

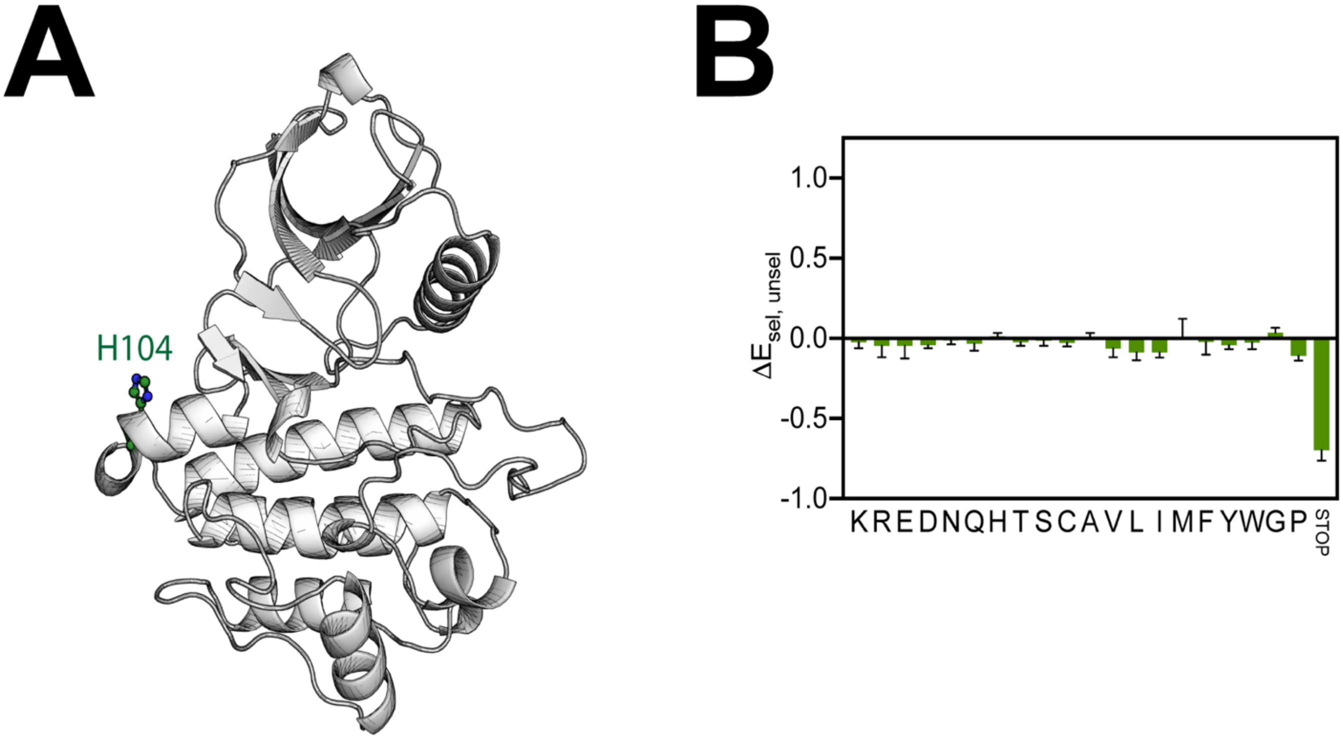


**Figure SI 6.** *Many residues are robust to mutation* **A.** H104 highlighted on the homology model of AncSZ. This residue is on the surface and the side chain is not involved in maintaining the structure or directly linked to catalysis. **B.** The average enrichment scores for substitutions to H104 in the bacterial two-hybrid (n=3). Most mutations have no effect on the enrichment score. However, a stop codon is a loss-of-function.

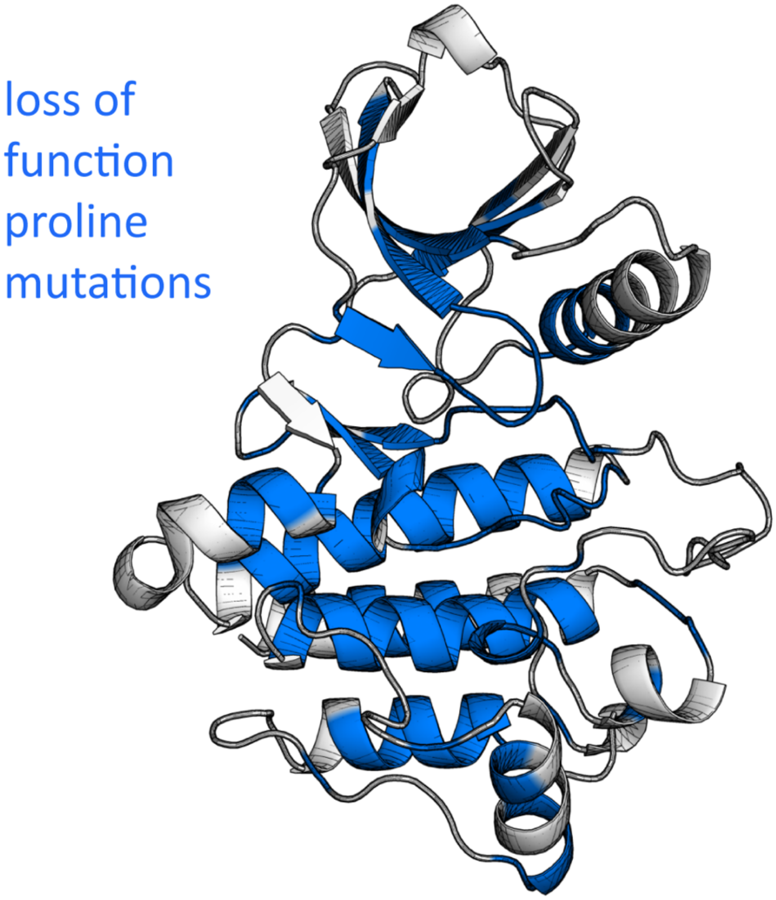


**Figure SI 7.** *Proline mutations are loss function in residues involved in the secondary structure.* Residues in which a proline substitution resulted in a loss of function in the bacterial two-hybrid assay are colored blue on the homology model of AncSZ. Many of these detrimental mutations occur in residues participating in secondary structure.
